# Striatal Direct Pathway Targets Npas1^+^ Pallidal Neurons

**DOI:** 10.1101/2020.09.02.273615

**Authors:** Qiaoling Cui, Xixun Du, Isaac Y. M. Chang, Arin Pamukcu, Varoth Lilascharoen, Brianna L. Berceau, Daniela García, Darius Hong, Uree Chon, Ahana Narayanan, Yongsoo Kim, Byung Kook Lim, C. Savio Chan

## Abstract

The classic basal ganglia circuit model asserts a complete segregation of the two striatal output pathways. Empirical data argue that, in addition to indirect-pathway striatal projection neurons (iSPNs), direct-pathway striatal projection neurons (dSPNs) innervate the external globus pallidus (GPe). However, the functions of the latter were not known. In this study, we interrogated the organization principles of striatopallidal projections and their roles in full-body movement in mice (both males and females). In contrast to the canonical motor-promoting response of dSPNs in the dorsomedial striatum (^DMS^dSPNs), optogenetic stimulation of dSPNs in the dorsolateral striatum (^DLS^dSPNs) suppressed locomotion. Circuit analyses revealed that dSPNs selectively target Npas1^+^ neurons in the GPe. In a chronic 6-hydroxydopamine lesion model of Parkinson’s disease, the dSPN-Npas1^+^ projection was dramatically strengthened. As ^DLS^dSPN-Npas1^+^ projection suppresses movement, the enhancement of this projection represents a circuit mechanism for the hypokinetic symptoms of Parkinson’s disease that has not been previously considered. In sum, our results suggest that dSPN input to the GPe is a critical circuit component that is involved in the regulation of movement in both healthy and parkinsonian states.

**Significance statement:** In the classic basal ganglia model, the striatum is described as a divergent structure—it controls motor and adaptive functions through two segregated, opposing output streams. However, the experimental results that show the projection from direct-pathway neurons to the external pallidum have been largely ignored. Here, we showed that this striatopallidal sub-pathway targets a select subset of neurons in the external pallidum and is motor-suppressing. We found that this sub-pathway undergoes changes in a Parkinson’s disease model. In particular, our results suggest that the increase in strength of this sub-pathway contributes to the slowness or reduced movements observed in Parkinson’s disease.

## Introduction

The basal ganglia are a group of subcortical nuclei that are critically involved in action control. As the primary input station of the basal ganglia macrocircuit, the dorsal striatum (dStr) computes an array of sensorimotor, cognitive, and motivational information. In addition, it is capable of supporting a wide repertoire of innate behaviors, such as feeding, grooming, and locomotion (Albin et al., 1995; Mink, 1996; Graybiel, 2008; Redgrave et al., 2010; Turner and Desmurget, 2010; Costa, 2011; Gerfen and Surmeier, 2011; Kravitz et al., 2012; Markowitz et al., 2018; Park et al., 2020; Weglage et al., 2020).

The dStr is composed of molecularly, synaptically, and functionally distinct subdivisions (Flaherty and Graybiel, 1994; Smith et al., 2004; Darvas and Palmiter, 2009, 2010; Pan et al., 2010; Nambu, 2011; Hintiryan et al., 2016; Hooks et al., 2018; Poulin et al., 2018; Martin et al., 2019; Alegre-Cortés et al., 2020; Ortiz et al., 2020). The dorsomedial striatum (DMS) and dorsolateral striatum (DLS) are thought to be involved in regulating goal-directed and habitual behavior, respectively (Yin and Knowlton, 2006; Balleine and O’Doherty, 2010; Redgrave et al., 2010; Cox and Witten, 2019). Although the role of DMS in locomotion is consistently observed across studies (Kravitz et al., 2010; Durieux et al., 2012; Cui et al., 2013; Freeze et al., 2013; Cazorla et al., 2014), the findings are counterintuitive, as this region of the dStr receives primarily non-motor cortical input (McGeorge and Faull, 1989; Flaherty and Graybiel, 1994; Znamenskiy and Zador, 2013; Oh et al., 2014; Hintiryan et al., 2016; Hunnicutt et al., 2016; Hooks et al., 2018; Chon et al., 2019). On the other hand, while the DLS receives inputs from premotor and motor regions, recent studies argue that it is not involved in locomotion but is required for gradual motor skill acquisition (Yin and Knowlton, 2006; Nambu, 2011; Durieux et al., 2012; Rothwell et al., 2014; O’Hare et al., 2016; Malvaez and Wassum, 2018). As different approaches are used across studies, the role of SPNs from the two spatial domains in controlling motor behaviors await further investigations.

While theories about how the dStr operates have been proposed, the precise cellular and circuit mechanisms involved remain to be determined. It has been difficult to disentangle the relative roles of the dStr and other nodes within the circuit in mediating these functions; this is in part because of the complexity of the cellular composition and circuit architecture within the basal ganglia. The classic circuit model asserts a complete segregation of the two striatal output pathways. Direct-pathway striatal projection neurons (dSPNs) facilitate movement via their projection to the substantia nigra pars reticulata; indirect-pathway striatal projection neurons (iSPNs) suppress movement via their projection to the external globus pallidus (GPe) (Albin et al., 1989; Alexander and Crutcher, 1990; Gerfen, 1992; DeLong and Wichmann, 2007; Kravitz et al., 2010). On the other hand, empirical data argue that, in addition to iSPNs, dSPNs innervate the GPe (Kawaguchi et al., 1990; Wu et al., 2000; Levesque and Parent, 2005; Fujiyama et al., 2011; Okamoto et al., 2020). Previous investigations suggest that the dSPN-GPe pathway shapes motor output (Cazorla et al., 2014). However, we only have limited information about the properties of this pathway. As the GPe contains a heterogenous population of neurons (Hernandez et al., 2015; Hegeman et al., 2016; Saunders et al., 2018; Abecassis et al., 2020; Cherian et al., 2020), the identity of neurons that receive dSPN input remains elusive. Here, we hypothesized that dSPNs regulate motor output by targeting a select subset of these neurons. By using transgenic and molecular tools, we dissected the cellular and spatial organization of the striatopallidal system and its implications in the regulation of locomotion.

## Methods

### Mice

All experiments detailed are in accordance with the Northwestern University Animal Care and Use Committee, the Institutional Animal Care and Use Committee at the University of California, San Diego, and are in compliance with the NIH Guide for the Care and Use of Laboratory Animals. *Adora2a*^Cre^ and *Drd1a*^Cre^ were obtained from MMRRC (Gong et al., 2007). *Drd1a*^tdTomato^ (Ade et al., 2011), *Pvalb*^tdTomato^ (Kaiser et al., 2016), *Pvalb*^Cre^ (Hippenmeyer et al., 2005), *R26*^LSL-tdTomato^ (Madisen et al., 2010), *R26*^FSF-LSL-tdTomato^ (Madisen et al., 2015), and *Tac1*^Cre^ (Harris et al., 2014) were obtained from Jackson Laboratory. *Npas1*^Cre-2A-tdTomato^ (Jax 027718, hereafter referred to as *Npas1*^Cre^) was generated in-house (Hernandez et al., 2015). Mice for all experiments were maintained on a C57BL/6J (Jax 000664) background. Mice were group-housed in standard cages on a 12-hr light-dark cycle. Both male and female mice were used in this study. To minimize the potential alteration of the phenotypes in mice carrying the transgene alleles, only hemizygous or heterozygous mice were used.

### Virus and tracer injections

Stereotaxic injections were performed as previously described (Cui et al., 2016). In brief, mice were anesthetized with isoflurane and immobilized on a stereotaxic frame (David Kopf Instruments). A small craniotomy (∼1 mm diameter) was made with a dental drill (Osada) for injection into a specific brain region. A virus or tracer was injected using a calibrated glass micropipette (VWR) at a rate of 0.3–0.5 μl/min. The micropipette was left *in situ* for 5–10 min post-injection to maximize tissue retention of virus or tracer and decrease capillary spread upon pipette withdrawal.

For optogenetic stimulation of direct-pathway striatal projection neurons (dSPNs) or indirect-pathway striatal projection neurons (iSPNs) in the dorsomedial striatum (DMS) or dorsolateral striatum (DLS) used for *ex vivo* experiments, a total 360 nl of EF1α-CreOn-hChR2(H134R)-eYFP adeno-associated virus (AAV) was injected unilaterally into the DMS (in mm: 0.9 rostral, 1.4 lateral, 3.4 and 3.0 ventral from bregma) or the DLS (in mm: 0.7 rostral, 2.3 lateral, 3.4 and 3.0 ventral from bregma) of *Drd1a*^Cre^ or *Adora2a*^Cre^ mice. For *in vivo* optogenetic stimulation of SPN soma or terminal fields, EF1α-CreOn-hChR2(H134R)-eYFP AAV was injected bilaterally into the DMS or DLS (540–720 nl per hemisphere). For control mice, EF1α-CreOn-eYFP AAV was injected into the same region with the same viral volume. For *in vivo* GtACR2 inhibition of SPN soma, hSyn1-SIO-stGtACR2-FusionRed AAV1 was injected bilaterally into the DMS or DLS (720 nl per hemisphere). To visualize the projection targets of dSPNs or iSPNs, hSyn-Flex-mRuby2-T2A-Synaptophysin-eGFP AAV (360 nl) was unilaterally injected into the DMS or DLS of *Drd1a*^Cre^, *Tac1*^Cre^, or *Adora2a*^Cre^ mice.

To determine whole-brain projections to the GPe, 90 nl of Cre-expressing lentivirus (LVretro-Cre) (Knowland et al., 2017) was mixed with cholera-toxin B subunit (CTb) conjugated to Alexa 488 (Thermo Fisher Scientific) at 1:1 ratio and was injected into the GPe (in mm: 0.2–0.25 caudal, 2.1–2.2 lateral, 4.1 ventral from bregma) of *R26*^LSL-tdTomato^ mice. For rabies virus (RbV)-based retrograde tracing, 200 nl of CreOn-mRuby2-TVA-RVG AAV (Shin et al., 2018) was first injected unilaterally into the GPe (in mm: 0.35 caudal and 2.0 lateral from bregma, 3.5 ventral from dura) of *Pvalb*^Cre^ and *Npas1*^Cre^ mice followed by EnvA-RVΔG-eGFP into the GPe three weeks later. To determine if single striatal neurons innervate both the GPe and substantia nigra pars reticulata (SNr), 90 nl of CTb 488 and 180 nl of Alexa 647-conjugated CTb (CTb 647; Thermo Fisher Scientific) were injected into the GPe (in mm: 0.2–0.25 caudal, 2.1–2.2 lateral, 4.1 ventral from bregma) and the SNr (in mm: 2.65 caudal, 1.5 lateral, 4.65 ventral from bregma, and 3.0 caudal, 1.44 lateral, 4.5 ventral from bregma), respectively, of *Drd1a*^tdTomato^ mice or *Tac1*^Cre^ mice to examine the colocalization of two CTbs with tdTomato or Cre. Alternatively, a CreOn-Flp canine virus (CAV) and CTb 647 were injected into the SNr together with hSyn-CreOn-FlpOn-ChR2-eYFP AAV into the dStr of *Drd1a*^Cre^ mice. To determine the topographical organization of the GPe to dStr projection, 360 nl of CTb 647 was injected into the DMS or DLS of C57BL/6J mice.

The locations of the targeted injections were visually inspected under epifluorescence microscopy in *ex vivo* slices or histologically verified *post hoc*. AAVs, LV, CAV, and RbV were used for this study. AAVs and LV were injected into mice (postnatal day 28–35, and postnatal day 55, respectively) at a final titer of 10^12^–10^13^ genome copies/ml, using the same standard procedures as stereotaxic injections (see above). CTbs were injected into mice at postnatal day 55–70. Recordings and immunohistological analyses were performed 28–40 days postoperatively. Fiber optic implantations were performed 21–35 days after viral injection. Fluorescence *in situ* hybridization was performed 1–2 weeks after the injection of RbV.

### Chronic 6-OHDA lesion

Unilateral lesion of the nigrostriatal system was produced by 6-hydroxydopamine HCl (6-OHDA) injection into the medial forebrain bundle (MFB) at postnatal day 28–35, as described previously (Cui et al., 2016). In brief, 6-OHDA (2.5–3 μg/μl) was dissolved in 0.9% (wt/vol) NaCl with 0.1% (wt/vol) ascorbic acid. Using identical procedures to those for stereotaxic injection of virus and tracer, 1 μl of 6-OHDA was injected into the MFB at (in mm) 0.70 caudal, 1.10 lateral, and 4.95 ventral from bregma. The extent of dopamine depletion was assessed using the cylinder test to quantify impairment in forelimb usage. Behavioral tests were carried out 3–5 weeks after 6-OHDA lesion. Contacts made by each forepaw on the wall of a clear glass cylinder (9 cm diameter) during spontaneous exploratory behavior were counted during a five-minute period. The asymmetry of the forelimb usage was defined as independent contralateral paw placement relative to that of the ipsilateral (to the injection) paw against the walls of the chamber during rearing and vertical or lateral explorations. Mice with less than 20% contralateral paw touches were deemed with severe dopamine loss and used for subsequent experiments. Electrophysiological experiments or immunohistological analyses were performed 4–6 weeks after 6-OHDA injection.

### Fiber implantation and behavior testing

Surgical procedures were the same as those for stereotaxic injections (see above). Fiber optic cannulae (250 μm core diameter, 0.66 NA) (Prizmatix) were bilaterally implanted into the target regions: the DMS (in mm: 0.9 rostral, 1.4 lateral, 3.0 ventral from bregma), the DLS (in mm: 0.7 rostral, 2.3 lateral, 3.0 ventral from bregma), the GPe (in mm: 0.3 caudal, 2.1 lateral, 3.7 ventral from bregma), or the SNr (in mm: 2.7 caudal, 1.4 lateral, 4.4 ventral from bregma). The fiber optic cannulae had a maximal output power of 12–18 mW measured at the tip. Implants were secured to the skull with dental cement (Parkell). Mice were allowed to recover for 1–2 weeks before behavioral testing.

Motor behavior induced by optogenetic stimulation was assessed in an open field. Behavioral testing was performed between 3:00 P.M. and 8:00 P.M. On the first day, the mice were allowed to freely explore the open field area (28 cm × 28 cm) for 25 min. On the second day, the implanted fiber-optic cannulae were connected to a 470 nm LED (Prizmatix). Five minutes after the mouse was placed in the open field arena, light stimulus trains (5 ms pulses at 10 Hz for 10 s) were delivered every min. A total of 20 stimulus trains were given to each mouse. Mice were videotaped with an overhead camera. The central position of each mouse was tracked with ETHOVISION (Noldus). Data for distance traveled over time were extracted. The locations of the targeted implantations were histologically verified *post hoc*.

### Behavioral tracking and classification

DeepLabCut (https://github.com/DeepLabCut/) (Mathis et al., 2018; Nath et al., 2019) was used for tracking body parts of mice in an open field arena. Eight body parts including the nose, ears, body center, side laterals (hip-joints), tail base, and tail end were labeled in top-down view videos. To create the training dataset, 1,674 distinct frames from 50 video recordings of open-field behavior were manually annotated. We used MobileNetV2-1-based network (Mathis et al., 2019; Sandler et al., 2019) with default parameters. The network was trained and refined for five rounds using default multi-step learning rates. Each round consists of 240,000–1,000,000 iterations, and the default multi-step learning rates were used. This trained network has a test error of 1.13 pixels and a training error of 4.82 pixels. Predictions of X-Y coordinates were processed using a median filter with a rolling window of five frames before further analysis. This network was then used to analyze all videos in this study.

To categorize motor behavior, DeepLabCut tracking data were first calibrated; the pixel-to-cm conversion for each video was determined by comparing the width of the arena in pixels to the actual width of the arena (28 cm). Based on the calibrated X-Y coordinates of labeled body parts, a set of movement metrics was generated for each frame. Mouse speed was measured as the body center speed. Mouse width was measured as the euclidean distance between the side laterals, and mouse length was measured as the euclidean distance between the nose and the tail base. Locomotion was defined as frames when the body center had a speed > 0.5 cm/s; motionless was defined as frames when the ears, body center, laterals, and tail base all had a speed ≤ 0.5 cm/s. To classify rearing, we constructed a random forest classifier in SimBA. 19,058 rearing frames from 35 video recordings of open-field behavior were extracted and manually annotated as rearing by three independent annotators. Supported and unsupported rearing behaviors were not differentiated. The start frame was defined as the frame in which the mouse lifted its forelimbs off the floor and extended its head upwards; the end frame was defined as the frame before the forelimbs made contact with the floor. The model was built with the following settings: n_estimators = 2,500, RF_criterion = entropy, RF_max_features = sqrt, RF_min_sample leaf = 2, and no oversampling or undersampling. 20% of the video frames were used for testing and the other 80% were used for training. The resulting classifier has a F1-score = 0.71, precision = 0.68, and recall = 0.74. The performance of this classifier was on par with those reported recently {049452}. The discrimination threshold was set at Pr = 0.31, and each instance of rearing had a minimum duration of 300 ms. Lastly, fine movement was defined as frames that did not fall into any of the categories mentioned above (i.e., locomotion, motionless, or rearing). Finally, example videos and the trained model are available on Github (https://github.com/saviochan/SimBA-OpenFieldArena) and Zenodo (https://zenodo.org/record/3964701#.XyB8yJ5KhPZ). The data generated by the analysis pipeline were processed using custom Python scripts. Codes are available online (https://github.com/saviochan/Python-Scripts/tree/master/OpenFieldArena_Behavior). Twenty-five different movement metrics were tracked. Event frequency, duration, and percent time spent were logged. ‘Light-period’ corresponds to 10 s of light delivery. ‘Pre-period’ and ‘post-period’ correspond to the 10 s epoch before and after light delivery, respectively. Fold changes were calculated by dividing the movement metric during light-period by that in pre-period.

To assess the relationship between the measured movement metrics, a correlation matrix was constructed from binned, time-series data. Rearing-motionless switch frequency and motionless-rearing switch frequency were excluded because of the low occurrence of events. Hierarchical clustering of movement metrics was performed in ClustVis (https://biit.cs.ut.ee/clustvis/) (Metsalu and Vilo, 2015). Twenty-five movement metrics were included in the analysis. Mice with targeted optogenetic stimulation of ^DMS^dSPNs, ^DMS^iSPNs, ^DLS^dSPNs and ^DLS^iSPNs, were included. Both rows and columns were clustered using correlation distance and average linkage. Movement metrics were centered and scaled.

### Fluorescence *in situ* hybridization

Brains were rapidly extracted and flash-frozen with isopentane chilled with dry ice in 70% ethanol. Coronal brain sections (25 μm) containing the dStr and the GPe were prepared on a cryostat (Leica). Brain sections were mounted directly onto glass slides and stored at −80 °C until further processed. Fluorescence *in situ* hybridization was conducted using commercial probes (Advanced Cell Diagnostics). Slides were fixed in 4% PFA for 15 min at 4 °C and subsequently dehydrated for 5–10 min with a series of ethanol at room temperature. Sections were then incubated with a protease pretreat-IV solution for 30 min, and washed with phosphate-buffered saline (PBS), before being incubated with probes for 2 h at 40 °C. After washes, the signal was amplified by incubating tissue sections in amplification buffers at 40 °C. After the final rinse, DAPI solution was applied to the sections. Slides were coverslipped and visualized with a confocal microscope (Olympus).

### Immunolabeling

Mice aged postnatal day 55–80 were anesthetized deeply with a ketamine-xylazine mixture and perfused transcardially first with 0.01 M PBS followed by fixative containing 4% (wt/vol) paraformaldehyde in PBS, pH 7.4. Brain tissue was then postfixed in the same fixative for 1–2 h at 4 °C. Tissue blocks containing the dStr and the GPe were sectioned using a vibrating microtome (Leica Instrument) at a thickness of 60 μm. Floating sections were blocked with 10% (vol/vol) normal goat or donkey serum (Gibco) and 0.1% (vol/vol) Triton X-100 in PBS for 30–60 min at room temperature and subsequently incubated with primary antibodies (**Table 1**) in the same solution for ∼24 h at 4 °C. After washes in PBS, the sections were incubated with Alexa-conjugated IgG secondary antibodies at 1:500 (Life Technologies) for 2 h at room temperature. The sections were then washed, mounted, and coverslipped. Immunoreactivity was examined on a laser-scanning confocal microscope with a 10× 0.45 numerical aperture (NA) air objective and a 60× 1.35 NA oil-immersion objective (Olympus).

**Table 1.** Primary antibodies used in this study.

| Antigen | Host species <sup>a</sup> | Clonality | Source | Catalog # | Working concentration | Dilution |
| --- | --- | --- | --- | --- | --- | --- |
| Cre | Gp | Polyclonal | Synaptic Systems | 257004 |  | 1:2,000 |
| RFP | Rb | Polyclonal | Clontech | 632496 | 1 µg/ml | 1:500 |
| Enkephalin | Rb | Polyclonal | Immunostar | 20065 |  | 1:200 |
| Gephyrin | Ms | Monoclonal | Synaptic Systems | 147011 | 2 µg/ml | 1:500 |
| GFP | Ck | Polyclonal | Abcam | ab13970 | 10 µg/ml | 1:1,000 |
| Npas1 | Gp | Polyclonal | Chan Laboratory<br>(Hernandez et al.,<br>2015) |  |  | 1:5,000 |
| Substance P | Rt | Monoclonal | Abcam | ab7340 |  | 1:200 |
| Substance P | Gp | Polyclonal | Abcam | ab10353 |  | 1:200 |
| tdTomato | Rt | Monoclonal | Kerafast | EST203 | 3.5 µg/ml | 1:500 |
| VGAT | Gp | Polyclonal | Synaptic Systems | 131004 |  | 1:500 |

Notes:
<sup>a</sup> Abbreviations: chicken (Ck), guinea pig (Gp), mouse (Ms), rabbit (Rb), and rat (Rt).

### Fiber density and synaptic contact quantifications

To measure fiber density of dSPNs in the dStr, GPe, and SNr, *Drd1a*^Cre^ mice were injected with EF1α-CreOn-hChR2(H134R)-eYFP AAV into the dStr. 60 μm-thick sections from lateral, intermediate, and medial levels (approximately 2.5 mm, 2.1 mm, 1.7 mm lateral from bregma) were sampled. eYFP fluorescence was enhanced with an antibody (**Table 1**). 10× images of eYFP fluorescence in the dStr, GPe, and SNr were captured with a confocal microscope. Fiber density was estimated based on eYFP fluorescence. Briefly, a region that encompassed the entirety of the dStr, GPe, or SNr with eYFP fluorescence was selected. Lower and upper threshold values were set using a thresholding function in Fiji (Schindelin et al., 2012). Integrated density was then measured.

To visualize dSPN terminals within the GPe, the same tissue sections as those used for fiber density analysis were used. These sections were co-stained with antibodies for gephyrin and vesicular GABA transporter (VGAT) (**Table 1**). In each section, three z-stacks spanning across the entire rostrocaudal axis of the GPe were imaged with a confocal microscope (Olympus) using a 60× objective with a 2× digital zoom. Five consecutive optical sections (1 µm interval) were captured for each z-stack. Quantification of putative synaptic contacts was performed with Fiji. The spatial relationship between labeled structures was examined across all three orthogonal planes. Synaptic contacts were deemed to be *bona fide* only if eYFP^+^ fibers were in close apposition with gephyrin^+^ or VGAT^+^ structures in all *x*-(≤ 0.2 μm), *y*-(≤ 0.2 μm), and *z*-(≤ 1 μm) planes.

To compare the density of terminals from dSPNs and iSPNs, tissue sections from *Drd1a*^Cre^, *Tac1*^Cre^, or *Adora2a*^Cre^ mice injected with hSyn-Flex-mRuby2-T2A-Synaptophysin-eGFP AAV into the DLS or DMS were used. eGFP fluorescence was enhanced with an antibody (**Table 1**). 60× z-stack images of eGFP fluorescence in the GPe were captured with a confocal microscope. Two regions of interest (ROIs) from dorsorostral and ventrocaudal territories of the GPe were imaged for each section. Integrated density was quantified from the maximal projection image of ten optical planes using the same thresholding function as described for fiber density analysis.

### Serial two-photon tomography

Serial two-photon tomography was used to map inputs to the GPe from the entire brain. Imaging and analysis were performed as previously described (Kim et al., 2017; Abecassis et al., 2020). Two weeks after LVretro-Cre and CTb-488 injection, mouse brains were fixed as described above. Brains were then transferred to PBS and stored at 4 °C until imaged. Brains were embedded in 4% agarose in 0.05 M phosphate buffer and cross-linked in 0.2% sodium borohydrate solution (in sodium borate buffer, pH 9.0–9.5). Each brain was imaged with a high-speed two-photon microscope with an integrated vibratome (TissueVision) at 1 μm at both *x–y* resolution with 280 *z*-sections in every 50 μm. A 910 nm two-photon laser (Coherent Technologies) was used for CTb488 and tdTomato excitation. A dichroic mirror (Chroma) and band-pass filters (Semrock) were used to separate green and red fluorescence signals. Emission signals were detected by GaAsP photomultiplier tubes (Hamamatsu). An automated, whole-brain cell counting and registration of the detected signal on a reference brain was applied as described before (Kim et al., 2017). The number of tdTomato neurons from each brain region was charted. The relative size of the input to the GPe was calculated by normalizing the total number of tdTomato neurons in the entire brain of each sample.

### Visualized recording in *ex vivo* brain slices

Mice aged postnatal day 55–75 were anesthetized with a ketamine-xylazine mixture and perfused transcardially with ice-cold artificial cerebrospinal fluid (aCSF) containing (in mM): 125 NaCl, 2.5 KCl, 1.25 NaH_2_PO4, 2.0 CaCl_2_, 1.0 MgCl_2_, 25 NaHCO_3_, and 12.5 glucose, bubbled continuously with carbogen (95% O_2_ and 5% CO_2_). The brains were rapidly removed, glued to the stage of a vibrating microtome (Leica Instrument), and immersed in ice-cold aCSF. Parasagittal (for DLS-GPe) or sagittal (for DMS-GPe) *ex vivo* slices containing the dStr and the GPe were cut at a thickness of 240 μm and transferred to a holding chamber, where they were submerged in aCSF at 35 °C for 30 min, and returned to room temperature before recording.

*Ex vivo* slices were then transferred to a small volume (∼0.5 ml) Delrin recording chamber that was mounted on a fixed-stage, upright microscope (Olympus). Neurons were visualized using differential interference contrast optics (Olympus), illuminated at 735 nm (Thorlabs), imaged with a 60× 1.0 NA water-immersion objective (Olympus) and a CCD camera (QImaging). PV^+^, PV^−^, Npas1^+^, and Npas1^−^ GPe neurons were identified by the presence or absence of somatic tdTomato fluorescence examined under epifluorescence microscopy with a daylight (6,500 K) LED (Thorlabs) and an appropriate filter cube (Semrock) from *Pvalb*^tdTomato^ and *Npas1*^Cre^ transgenic mice.

Recordings were made at room temperature (20–22 °C) with patch electrodes fabricated from capillary glass (Sutter Instruments) pulled on a Flaming-Brown puller (Sutter Instruments) and fire-polished with a microforge (Narishige) immediately before use. Pipette resistance was typically ∼3–5 MΩ. Internal solution for voltage-clamp recordings of inhibitory postsynaptic currents contained (in mM): 120 CsCl, 10 Na_2_phosphocreatine, 5 HEPES, 5 tetraethylammonium-Cl, 2 Mg_2_ATP, 1 QX314-Cl, 0.5 Na_3_GTP, 0.5 CaCl_2_, 0.25 EGTA, and 0.2% (wt/vol) biocytin, pH adjusted to 7.25–7.30 with CsOH. This internal solution had an osmolarity of ∼290 mOsm. Somatic whole-cell patch-clamp recordings were obtained with an amplifier (Molecular Devices). The signal for voltage-clamp recordings was filtered at 1 kHz and digitized at 10 kHz with a digitizer (Molecular Devices). Stimulus generation and data acquisition were performed using pClamp (Molecular Devices).

For optogenetic experiments, blue (peak at ∼450 nm) excitation wavelength from two daylight (6,500 K) LEDs (Thorlabs) was delivered to the tissue slice bidirectionally from both the 60× water immersion objective and the 0.9 NA air condenser with the aid of 520 nm dichroic beamsplitters (Semrock). The field of illumination that centered around the recorded cells was ∼500 μm in diameter. The duration for all light pulses was 2 ms. All recordings were made in the presence of R-CPP (10 μM) and NBQX (5 μM) to prevent the confounding effects of incidental stimulation of glutamatergic inputs. CGP55845 (1 μM) was also included to prevent GABA_B_ receptor-mediated modulation. In a subset of experiments, 2–4 mM Sr^2+^ was used to replace external Ca^2+^ to measure quantal events (Bekkers and Clements, 1999; Xu-Friedman and Regehr, 1999, 2000; McGarry and Carter, 2017).

Off-line data analyses were performed with ClampFit (Molecular Devices) and MiniAnalysis (Synaptosoft). Paired-pulse ratios were calculated by dividing the second inhibitory postsynaptic current (IPSC) amplitude by the first IPSC amplitude (Kim and Alger, 2001). For strontium-based quantal analysis, events within 300 ms after the striatal stimulation were quantified.

### Drugs

*R*-baclofen, *R*-CPP, LY341495, and NBQX disodium salt were obtained from Tocris. CGP55845, QX314-Cl, and SR95531 were obtained from Abcam. Na_3_GTP and tetrodotoxin were from Roche and Alomone Laboratories, respectively. Other reagents not listed above were from Sigma-Aldrich. Drugs were dissolved as stock solutions in either water or DMSO and aliquoted and frozen at –30 °C prior to use. Each drug was diluted to the appropriate concentrations by adding to the perfusate immediately before the experiment. The final concentration of DMSO in the perfusate was < 0.1%.

### Experimental design and statistical analyses

General graphing and statistical analyses were performed with MATLAB (MathWorks), Prism (GraphPad), JASP (https://jasp-stats.org), and the R environment (https://www.r-project.org). Custom analysis codes are available on GitHub (https://github.com/chanlab). Sample size (*n* value) is defined by the number of observations (i.e., ROIs, synaptic contacts, neurons, sections, or mice). When percentages are presented, *n* values represent only positive observations. No statistical method was used to predetermine sample size. Data in the main text are presented as median values ± median absolute deviations (MADs) (Leys et al., 2013) as measures of central tendency and statistical dispersion, respectively. Box plots are used for graphic representation of population data unless stated otherwise (Krzywinski and Altman, 2014; Streit and Gehlenborg, 2014; Nuzzo, 2016). The central line represents the median, the box edges represent the interquartile ranges, and the whiskers represent 10–90th percentiles. Normal distributions of data were not assumed. Individual data points were visualized for small sizes or to emphasize variability in the datasets. Non-parametric statistics were used throughout. Comparisons of unrelated samples were performed using a Mann–Whitney *U* test. The Wilcoxon signed rank test was used for pairwise comparisons for related samples. The Fisher’s exact test was used for categorical data. The Spearman exact test was used for evaluating the correlation between variables. Unless < 0.0001 or > 0.99, exact *P* values (two-tailed) are reported in the text. To avoid arbitrary cutoffs and visual clutter, levels of significance are not included in the figures.

## Results

### ^DLS^dSPNs are motor-suppressing

To study the behavioral roles of SPN subtypes from the DMS and DLS in motor regulation, we first selectively stimulated ^DMS^dSPNs and ^DMS^iSPNs using ChR2, an excitatory opsin (Boyden et al., 2005), as a proof of concept. *Drd1a*^Cre^ and *Adora2a*^Cre^ mice, two well-characterized, pathway-specific driver lines were used in conjunction with Cre-inducible adeno-associated viruses (AAVs) to confer transgene expression in dSPNs and iSPNs, respectively (Gong et al., 2007; Lobo et al., 2010; Gerfen et al., 2013; Okamoto et al., 2020). Motor behavior was monitored in an open-field arena (see Methods). Consistent with the findings from prior studies (Kravitz et al., 2010; Durieux et al., 2012; Freeze et al., 2013), stimulation of ^DMS^dSPNs and ^DMS^iSPNs led to canonical movement promotion and suppression, respectively, as measured by the change in speed (^DMS^dSPNs = +0.71 ± 0.28 fold, *n* = 9 mice, *P* = 0.0039; ^DMS^iSPNs = –0.43 ± 0.05 fold, *n* = 11 mice, *P* = 0.00098). On the contrary, optogenetic stimulation of ^DLS^dSPNs and ^DLS^iSPNs suppressed and promoted movement, respectively (^DLS^dSPNs = –0.34 ± 0.24 fold, *n* = 13 mice, *P* = 0.0017; ^DLS^iSPNs = +0.38 ± 0.31 fold, *n* = 15 mice, *P* = 0.00061) (**Figure 1a–d**). The motor effects induced with ChR2 activation were different from their corresponding eYFP controls (**Figure 1e & g**), arguing that the findings were not artifacts of light delivery (Owen et al., 2019).

**Figure 1.**
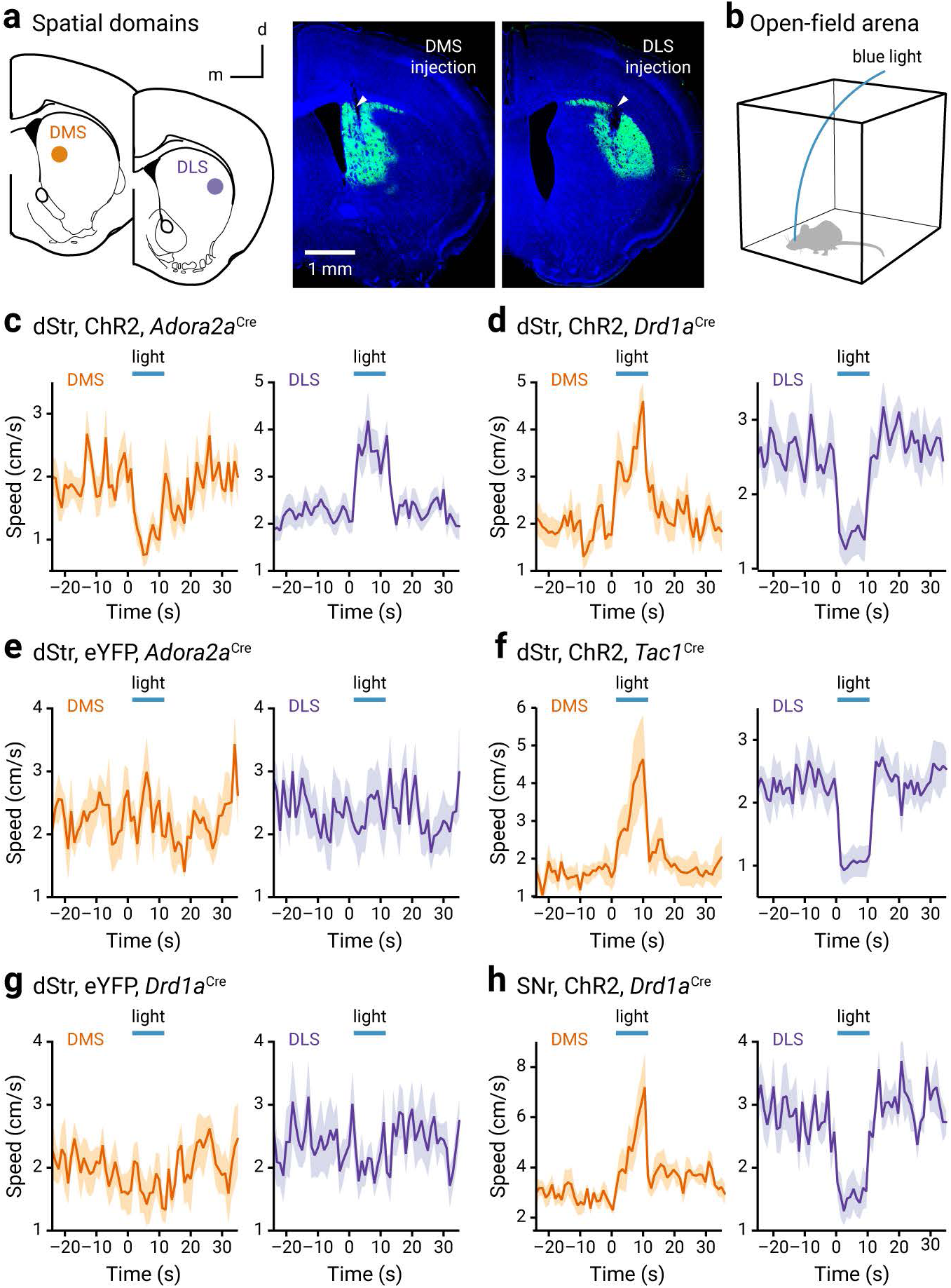
Optogenetic stimulation of SPN subtypes induces unique changes in locomotor speed. **a.** *Left*, a schematic diagram showing injection in the DMS and DLS. Cre-inducible (CreOn) ChR2-eYFP and eYFP AAV injections were targeted to the DMS (in mm: 0.9 rostral, 1.4 lateral, 3.4 and 3.0 ventral from bregma) or the DLS (in mm: 0.7 rostral, 2.3 lateral, 3.4 and 3.0 ventral from bregma) of *Adora2a*^Cre^, *Drd1a*^Cre^, or *Tac1*^Cre^ mice. *Right*, coronal brain sections showing the viral spread in the dorsal striatum. Green = ChR2-eYFP; blue = DAPI. Tracks produced by the fiber cannulae are marked by arrowheads. Scale bar applies to both images. **b.** An open-field arena (28 cm x 28 cm) was used for examining the locomotor activity of test subjects (see Methods). **c**–**g**. Changes in speeds are shown with light delivery (blue horizontal lines) in the DMS (orange) and DLS (purple) of *Adora2a*^Cre^, *Drd1a*^Cre^, and *Tac1*^Cre^ mice that expressed ChR2 or eYFP. **h**. Changes in speeds are shown with light delivery (blue horizontal lines) in the medial (left) and lateral (right) SNr of *Drd1a*^Cre^ mice that expressed ChR2. CreOn-ChR2-eYFP AAV injections were targeted to the DMS and the DLS, respectively.

While the validity of the *Adora2a*^Cre^ mouse has been unequivocally confirmed with the examination of the gross axonal projection pattern of Cre-expressing neurons, it is more challenging to confirm the absolute specificity of the *Drd1a*^Cre^ mouse because of the multi-projectional nature of dSPNs. As substance P (and its mRNA, *Tac1*) is selectively enriched in dSPNs (Gerfen and Young, 1988; Gerfen et al., 1990; Lobo et al., 2006; Heiman et al., 2008; Gokce et al., 2016), *Tac1*^Cre^ knock-in mice were employed to confirm the inferences drawn from *Drd1a*^Cre^ mice. Consistent with the observations in *Drd1a*^Cre^ mice, optogenetic stimulation of ^DLS^dSPNs in *Tac1*^Cre^ mice led to suppression of locomotor speed (^DLS^dSPNs = –0.61 ± 0.11 fold, *n* = 10 mice, *P* = 0.0020). This effect was different (*P* = 0.00067) from that induced by optogenetic stimulation of ^DMS^dSPNs in *Tac1*^Cre^ mice (^DMS^dSPNs = 0.69 ± 0.54 fold, *n* = 5 mice) (**Figure 1f**). By interrogating with optogenetic approaches, here we conclude that striatal spatial subdomains exhibit divergent locomotor regulation.

### High-dimensional analyses confirm diametric motor responses produced by SPN subtypes

To survey the full range of motor behaviors, a machine learning approach was used to track body kinematics and movement dynamics (Cherian et al., 2020). These data are summarized in a heatmap format (**Figure 2a**). By performing hierarchical clustering of movement metrics, we showed that the optogenetic stimulation of ^DMS^dSPNs and ^DMS^iSPNs induced congruent changes in motor behaviors across all mice examined—mice with targeted optogenetic stimulation of ^DMS^dSPNs and ^DMS^iSPNs fell into distinct clusters. On the contrary, mice with targeted optogenetic stimulation of ^DLS^dSPNs and ^DLS^iSPNs were intermixed with the two main clusters formed by mice that were targeted for optogenetic stimulation of ^DMS^dSPNs and ^DMS^iSPNs. The different motor patterns induced by selective stimulation of specific SPNs can be readily observed when ‘motionless’, ‘fine movement’, ‘rearing’, and ‘locomotion’ are charted (**Figure 2b & c**).

**Figure 2.**
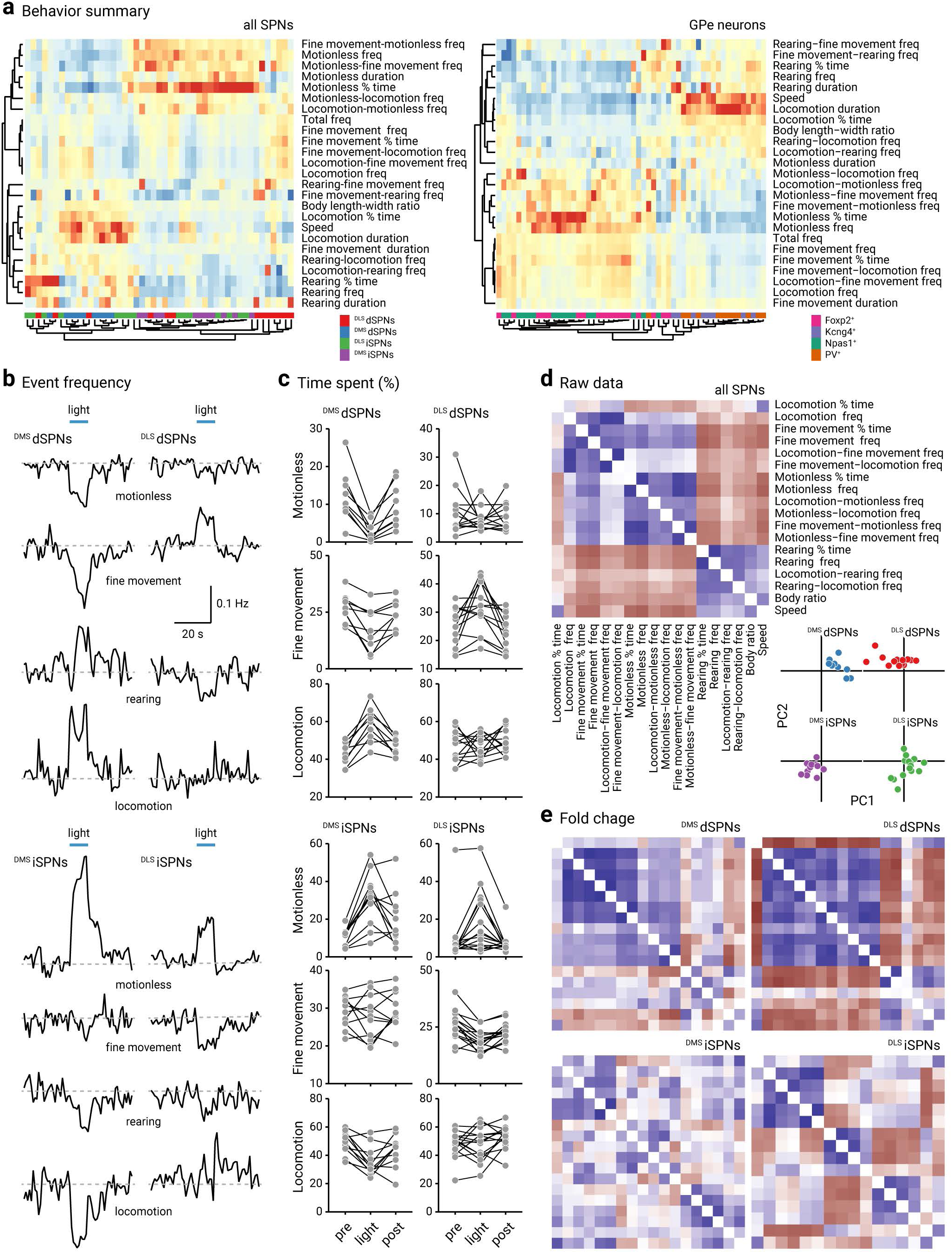
Optogenetic stimulation of SPN subtypes produces unique changes in movement metrics. **a.** *Left*, a heatmap summarizing motor responses of mice to optogenetic stimulation of ^DLS^dSPNs (red), ^DMS^dSPNs (blue), ^DLS^iSPNs (green), ^DMS^iSPNs (purple). Twenty-five movement metrics were measured to fully capture the behavioral structures. Each of the 25 rows represents the fold change of movement metrics. Warm colors (red) represent positive changes; cool colors (blue) represent negative changes. Rows and columns were sorted using hierarchical clustering. Dendrograms are divided into two main arms; metrics on the upper arm are positively correlated with ‘total frequency’, while metrics on the lower arm are negatively correlated with ‘total frequency’. Each column is a mouse; 48 mice were used in this and all subsequent analyses in **b**–**e** (^DLS^dSPNs = 13 mice, ^DMS^dSPNs = 9 mice, ^DLS^iSPNs = 15 mice, ^DMS^iSPNs = 11 mice). *Right*, a heatmap summarizing motor responses of mice to optogenetic stimulation of genetically-defined neurons in the GPe. The neurons of interest are Foxp2^+^ (pink), Kcng4^+^ (purple), Npas1^+^ (green), PV^+^ (orange). The plot was reproduced from Cherian et al. (Cherian et al., 2020). **b.** Mean changes in the event frequency of motionless, fine movement, rearing, and locomotion upon optogenetic stimulation (blue horizontal lines) of ^DMS^dSPNs, ^DLS^dSPNs, ^DMS^iSPNs, and ^DLS^iSPNs. Scale bar applies to all traces. **c.** Slopegraphs showing the fraction of time spent for motionless, fine movement, and locomotion in mice and the effect with optogenetic stimulation of selective neuron types. Each connected line represents a mouse. **d.** A correlation matrix constructed using data from *Adora2a*^Cre^ and *Drd1a*^Cre^ mice transduced with CreOn-ChR2-eYFP AAV. Eighteen parameters were included in this matrix. Blue colors indicate positive correlations, whereas brown colors indicate negative correlations. *Inset*, Principal component analysis plots showing the distribution of ^DMS^dSPNs (blue), ^DLS^dSPNs (red), ^DMS^iSPNs (purple), and ^DLS^iSPNs (green). Fold changes of twenty three movement metrics with optogenetic stimulation were used in this analysis. **e.** A correlation matrix constructed from fold changes in movement metrics following optogenetic stimulation of ^DMS^dSPNs, ^DLS^dSPNs, ^DMS^iSPNs, and ^DLS^iSPNs. Blue colors indicate positive correlations, brown colors indicate negative correlations.

Consistent with the well-established roles of ^DMS^dSPNs and ^DMS^iSPNs, optogenetic stimulation of ^DMS^dSPNs and ^DMS^iSPNs led to coordinated changes in motionless and locomotion. On the contrary, the changes induced by optogenetic stimulation of ^DLS^dSPNs and ^DLS^iSPNs resulted in less marked or coherent changes in motionless and locomotion; the changes in net motor output as measured by the distance traveled were primarily driven by the changes in movement speed. In addition, fine movement was increased with ^DLS^dSPNs stimulation and decreased with ^DLS^iSPNs stimulation. The differences in the movement dynamics induced with optogenetic stimulation of SPN subtypes can be found in **Table 2**. Lastly, the uniqueness of each SPN population in the induced motor behavior is more clearly illustrated in the correlation matrix and principal component analysis of movement metrics (**Figure 2d & e**).

**Table 2.** Summary of optogenetic effects on behavior metrics.

|  | DMS<sub>1</sub>dSPNs, Chr2 (n = 9) |  |  |  | DMS<sub>1</sub>iSPNs, Chr2 (n = 11) |  |  |  |
| --- | --- | --- | --- | --- | --- | --- | --- | --- |
|  | Pre | Light | Fold change | P value <sup>c</sup> | Pre | Light | Fold change | P value |
| Speed (cm/s) | 2.43 ± 0.60 | 4.12 ± 0.56 | 1.90 ± 0.25 | **0.0039** | 2.47 ± 0.43 | 1.23 ± 0.19 | 0.52 ± 0.09 | **0.00098** |
| Body length-width ratio | 2.34 ± 0.13 | 2.75 ± 0.15 | 1.14 ± 0.11 | **0.0039** | 2.32 ± 0.19 | 2.16 ± 0.12 | 0.97 ± 0.04 | 0.083 |
| Total frequency <sup>a</sup> | 35.33 ± 3.43 | 24.30 ± 8.03 | 0.76 ± 0.16 | **0.0039** | 36.40 ± 3.30 | 38.70 ± 2.70 | 1.12 ± 0.09 | 0.31 |
| Locomotion frequency | 13.20 ± 0.66 | 11.70 ± 2.63 | 0.91 ± 0.18 | 0.20 | 14.60 ± 0.80 | 13.30 ± 0.55 | 0.90 ± 0.04 | **0.0098** |
| Fine movement frequency | 15.44 ± 2.24 | 10.10 ± 4.57 | 0.82 ± 0.14 | **0.0039** | 15.00 ± 0.75 | 15.70 ± 1.70 | 1.04 ± 0.08 | 0.81 |
| Rearing frequency | 1.40 ± 0.70 | 1.10 ± 0.70 | 0.97 ± 0.32 | 0.97 | 1.10 ± 0.30 | 0.30 ± 0.20 | 0.29 ± 0.11 | **0.00098** |
| Motionless frequency | 5.40 ± 1.60 | 1.50 ± 1.10 | 0.33 ± 0.14 | **0.0039** | 5.00 ± 1.29 | 10.50 ± 1.75 | 1.88 ± 0.34 | **0.00098** |
| Locomotion % time | 42.80 ± 3.70 | 59.90 ± 6.10 | 1.37 ± 0.12 | **0.0039** | 49.60 ± 4.90 | 32.50 ± 4.80 | 0.66 ± 0.10 | **0.0029** |
| Fine movement % time | 26.78 ± 3.92 | 16.80 ± 7.87 | 0.79 ± 0.13 | **0.0039** | 28.93 ± 3.07 | 27.90 ± 5.60 | 1.03 ± 0.08 | 0.64 |
| Rearing % time | 15.70 ± 6.70 | 13.40 ± 8.40 | 1.11 ± 0.57 | 0.65 | 12.60 ± 5.90 | 3.70 ± 3.40 | 0.39 ± 0.18 | **0.00098** |
| Motionless % time | 10.20 ± 2.60 | 1.80 ± 1.65 | 0.18 ± 0.10 | **0.0039** | 12.50 ± 1.00 | 32.30 ± 5.40 | 2.84 ± 0.34 | **0.00098** |
| Locomotion duration <sup>b</sup> | 0.34 ± 0.03 | 0.60 ± 0.14 | 1.70 ± 0.41 | **0.0039** | 0.34 ± 0.05 | 0.34 ± 0.07 | 0.95 ± 0.23 | 0.21 |
| Fine movement duration | 0.18 ± 0.01 | 0.16 ± 0.01 | 0.93 ± 0.07 | **0.027** | 0.18 ± 0.01 | 0.17 ± 0.01 | 0.94 ± 0.05 | **0.019** |
| Rearing duration | 1.16 ± 0.07 | 1.01 ± 0.30 | 0.84 ± 0.29 | 0.82 | 0.96 ± 0.11 | 0.88 ± 0.35 | 1.28 ± 0.56 | 0.43 |
| Motionless duration | 0.20 ± 0.13 | 0.11 ± 0.01 | 0.61 ± 0.08 | **0.0039** | 0.19 ± 0.03 | 0.30 ± 0.07 | 1.67 ± 0.32 | **0.0020** |
| Locomotion-fine movement frequency | 10.70 ± 9.73 | 9.30 ± 8.33 | 1.01 ± 0.24 | 0.50 | 11.79 ± 11.24 | 9.10 ± 8.38 | 0.72 ± 0.13 | **0.0020** |
| Locomotion-rearing frequency | 1.50 ± 0.28 | 1.58 ± 0.18 | 1.14 ± 0.13 | 0.31 | 1.43 ± 0.32 | 1.00 ± 0.00 | 0.57 ± 0.29 | **0.031** |
| Locomotion-motionless frequency | 2.00 ± 0.43 | 1.20 ± 0.20 | 0.77 ± 0.06 | **0.0039** | 2.09 ± 0.42 | 2.89 ± 0.51 | 1.38 ± 0.35 | **0.0088** |
| Fine movement-locomotion frequency | 10.22 ± 0.63 | 9.20 ± 3.24 | 1.02 ± 0.30 | 0.48 | 11.70 ± 0.77 | 9.40 ± 2.65 | 0.80 ± 0.12 | **0.0020** |
| Fine movement-rearing frequency | 1.00 ± 0.17 | 1.00 ± 0.00 | 0.86 ± 0.46 | 0.44 | 1.00 ± 0.00 | 0.00 ± 0.00 | 0.00 ± 0.00 | > 0.99 |
| Fine movement-motionless frequency | 3.67 ± 1.13 | 2.00 ± 0.90 | 0.45 ± 0.11 | **0.0039** | 4.13 ± 0.75 | 6.75 ± 1.00 | 1.56 ± 0.36 | **0.00098** |
| Rearing-locomotion frequency | 1.67 ± 0.33 | 1.43 ± 0.29 | 1.07 ± 0.18 | 0.81 | 1.50 ± 0.13 | 1.00 ± 0.00 | 0.67 ± 0.17 | **0.0078** |
| Rearing-fine movement frequency | 1.00 ± 0.00 | 1.00 ± 0.00 | 1.00 ± 0.14 | 0.63 | 1.00 ± 0.33 | 1.00 ± 0.00 | 1.00 ± 0.33 | > 0.99 |
| Motionless-locomotion frequency | 2.17 ± 0.37 | 1.40 ± 0.40 | 0.71 ± 0.07 | **0.0078** | 2.17 ± 0.29 | 2.60 ± 0.35 | 1.42 ± 0.27 | **0.0059** |
| Motionless-fine movement frequency | 4.13 ± 1.03 | 1.75 ± 0.75 | 0.43 ± 0.11 | **0.0039** | 4.67 ± 1.17 | 6.29 ± 1.17 | 1.67 ± 0.35 | **0.00098** |

|  | DLS<sub>+</sub>dSPNs, Chr2 (n = 13) |  |  |  | DLS<sub>+</sub>iSPNs, Chr2 (n = 15) |  |  |  |
| --- | --- | --- | --- | --- | --- | --- | --- | --- |
|  | Pre | Light | Fold change | P value | Pre | Light | Fold change | P value |
| Speed (cm/s) | 3.04 ± 0.92 | 1.32 ± 0.44 | 0.59 ± 0.34 | **0.0061** | 2.75 ± 0.32 | 3.80 ± 0.65 | 1.35 ± 0.30 | **0.0012** |
| Body length-width ratio | 2.54 ± 0.14 | 2.46 ± 0.12 | 1.01 ± 0.06 | 0.84 | 2.39 ± 0.15 | 2.52 ± 0.11 | 1.06 ± 0.06 | **0.035** |
| Total frequency | 32.20 ± 6.40 | 37.80 ± 5.33 | 1.13 ± 0.15 | 0.057 | 35.20 ± 3.87 | 27.85 ± 3.25 | 0.85 ± 0.12 | **0.0081** |
| Locomotion frequency | 13.25 ± 1.42 | 15.53 ± 1.55 | 1.17 ± 0.11 | **0.0024** | 14.17 ± 2.28 | 10.22 ± 0.81 | 0.72 ± 0.11 | **0.00085** |
| Fine movement frequency | 13.92 ± 2.88 | 16.70 ± 2.10 | 1.18 ± 0.20 | **0.017** | 14.31 ± 1.61 | 11.15 ± 1.60 | 0.80 ± 0.12 | **0.00085** |
| Rearing frequency | 1.40 ± 0.32 | 0.70 ± 0.37 | 0.50 ± 0.31 | **0.0081** | 1.80 ± 0.45 | 1.17 ± 0.61 | 0.90 ± 0.43 | 0.61 |
| Motionless frequency | 3.70 ± 1.40 | 4.50 ± 0.75 | 1.16 ± 0.45 | 0.72 | 3.07 ± 0.90 | 5.62 ± 1.85 | 1.45 ± 0.55 | **0.027** |
| Locomotion % time | 48.40 ± 7.47 | 45.00 ± 5.00 | 1.03 ± 0.13 | 0.84 | 49.43 ± 4.07 | 49.92 ± 9.12 | 0.99 ± 0.16 | 0.92 |
| Fine movement % time | 24.83 ± 6.42 | 32.30 ± 8.40 | 1.17 ± 0.24 | **0.021** | 26.38 ± 4.62 | 18.77 ± 3.21 | 0.75 ± 0.08 | **0.00085** |
| Rearing % time | 17.30 ± 4.01 | 10.75 ± 6.55 | 0.78 ± 0.26 | 0.057 | 17.10 ± 7.56 | 11.77 ± 6.19 | 0.96 ± 0.53 | 0.36 |
| Motionless % time | 6.08 ± 3.42 | 6.70 ± 1.60 | 0.90 ± 0.51 | 0.49 | 4.79 ± 1.37 | 12.38 ± 8.28 | 2.01 ± 1.10 | **0.0051** |
| Locomotion duration | 0.38 ± 0.08 | 0.32 ± 0.07 | 0.86 ± 0.20 | 0.13 | 0.38 ± 0.05 | 0.51 ± 0.10 | 1.18 ± 0.20 | **0.00031** |
| Fine movement duration | 0.17 ± 0.01 | 0.18 ± 0.01 | 1.02 ± 0.10 | 0.22 | 0.17 ± 0.01 | 0.16 ± 0.01 | 0.91 ± 0.06 | 0.095 |
| Rearing duration | 1.21 ± 0.11 | 1.48 ± 0.28 | 1.10 ± 0.22 | 0.11 | 1.03 ± 0.23 | 0.98 ± 0.18 | 0.90 ± 0.32 | 0.22 |
| Motionless duration | 0.18 ± 0.02 | 0.13 ± 0.01 | 0.73 ± 0.14 | **0.00073** | 0.15 ± 0.02 | 0.24 ± 0.07 | 1.42 ± 0.43 | **0.0026** |
| Locomotion-fine movement frequency | 10.80 ± 9.72 | 12.58 ± 11.46 | 1.26 ± 0.11 | **0.00098** | 11.70 ± 10.96 | 6.46 ± 5.93 | 0.62 ± 0.12 | **0.00043** |
| Locomotion-rearing frequency | 1.50 ± 0.20 | 1.40 ± 0.40 | 0.97 ± 0.17 | 0.75 | 1.43 ± 0.26 | 1.33 ± 0.33 | 1.00 ± 0.25 | 0.81 |
| Locomotion-motionless frequency | 1.70 ± 0.27 | 2.14 ± 0.46 | 1.25 ± 0.31 | 0.13 | 2.00 ± 0.33 | 2.08 ± 0.50 | 1.08 ± 0.25 | 0.22 |
| Fine movement-locomotion frequency | 10.60 ± 1.15 | 12.90 ± 1.33 | 1.28 ± 0.17 | **0.00024** | 11.64 ± 1.94 | 6.38 ± 1.08 | 0.62 ± 0.12 | **0.00037** |
| Fine movement-rearing frequency | 1.00 ± 0.00 | 1.00 ± 0.00 | 0.88 ± 0.21 | 0.88 | 1.33 ± 0.33 | 1.00 ± 0.25 | 0.77 ± 0.23 | 0.07 |
| Fine movement-motionless frequency | 3.63 ± 1.10 | 4.00 ± 1.00 | 1.03 ± 0.41 | 0.79 | 2.80 ± 0.63 | 3.45 ± 1.31 | 1.42 ± 0.35 | **0.035** |
| Rearing-locomotion frequency | 1.33 ± 0.17 | 1.00 ± 0.00 | 0.89 ± 0.14 | 0.12 | 1.50 ± 0.35 | 1.43 ± 0.24 | 1.00 ± 0.25 | 0.78 |
| Rearing-fine movement frequency | 1.00 ± 0.00 | 1.00 ± 0.00 | 1.00 ± 0.20 | 0.81 | 1.00 ± 0.00 | 1.00 ± 0.00 | 1.00 ± 0.06 | 0.88 |
| Motionless-locomotion frequency | 2.15 ± 0.35 | 2.20 ± 0.20 | 1.04 ± 0.25 | 0.64 | 1.86 ± 0.36 | 2.00 ± 0.50 | 1.11 ± 0.16 | 0.2 |
| Motionless-fine movement frequency | 3.20 ± 0.76 | 4.38 ± 0.88 | 1.36 ± 0.68 | 0.3 | 2.75 ± 0.58 | 3.36 ± 1.30 | 1.39 ± 0.50 | 0.12 |

Notes:
<sup>a</sup> Unit for all frequencies is count/10 s.
<sup>b</sup> Unit for all durations is seconds.
<sup>c</sup> Wilcoxon test; two-tailed exact *P* values are shown. *P* < 0.05 are shown in bold.

To confirm the observed motor effects were indeed selectively associated with dSPNs from different spatial subdomains, we optogenetically stimulated their terminals in the substantia nigra pars reticulata (SNr). Consistent with the motor effect of intrastriatal stimulation of ^DLS^dSPNs, stimulation of their terminals in the lateral SNr led to movement suppression (–0.42 ± 0.14 fold, *n* = 7 mice, *P* = 0.016). As expected, this effect was opposite to the movement promotion induced with stimulation of ^DMS^dSPN terminals in the medial SNr (+0.68 ± 0.32 fold, *n* = 6 mice, *P* = 0.031) (**Figure 1h**). The motor effects induced with ChR2 activation were different from their corresponding eYFP controls (lateral SNr: *P* = 0.0025; medial SNr: *P* = 0.0095). The changes in motor patterns with stimulation of dSPN terminals in the SNr are shown in **Figure 3a**. Similar to the findings with stimulation of ^DLS^dSPNs and their terminals in the lateral SNr, optogenetic stimulation of their terminals within the GPe produced motor suppression (–0.34 ± 0.21 fold, *n* = 16 mice, *P* = 0.00021). The effect was different (*P* = 0.0046) from mice in which no ChR2 (i.e., eYFP only) was expressed (+0.01 ± 0.07 fold, *n* = 6 mice, *P* > 0.99). The fold changes were indistinguishable between stimulation of soma and their terminals in the GPe (*P* = 0.88). As ChR2 activation evoked action potentials that propagate both orthodromically and antidromically, it was difficult to pinpoint the precise effector loci responsible for the motor effects. Nonetheless, the *in vivo* optogenetic interrogation here collectively demonstrated the existence of parallel pathways that emerge from dStr spatial subdomains and their topographical organization.

**Figure 3.**
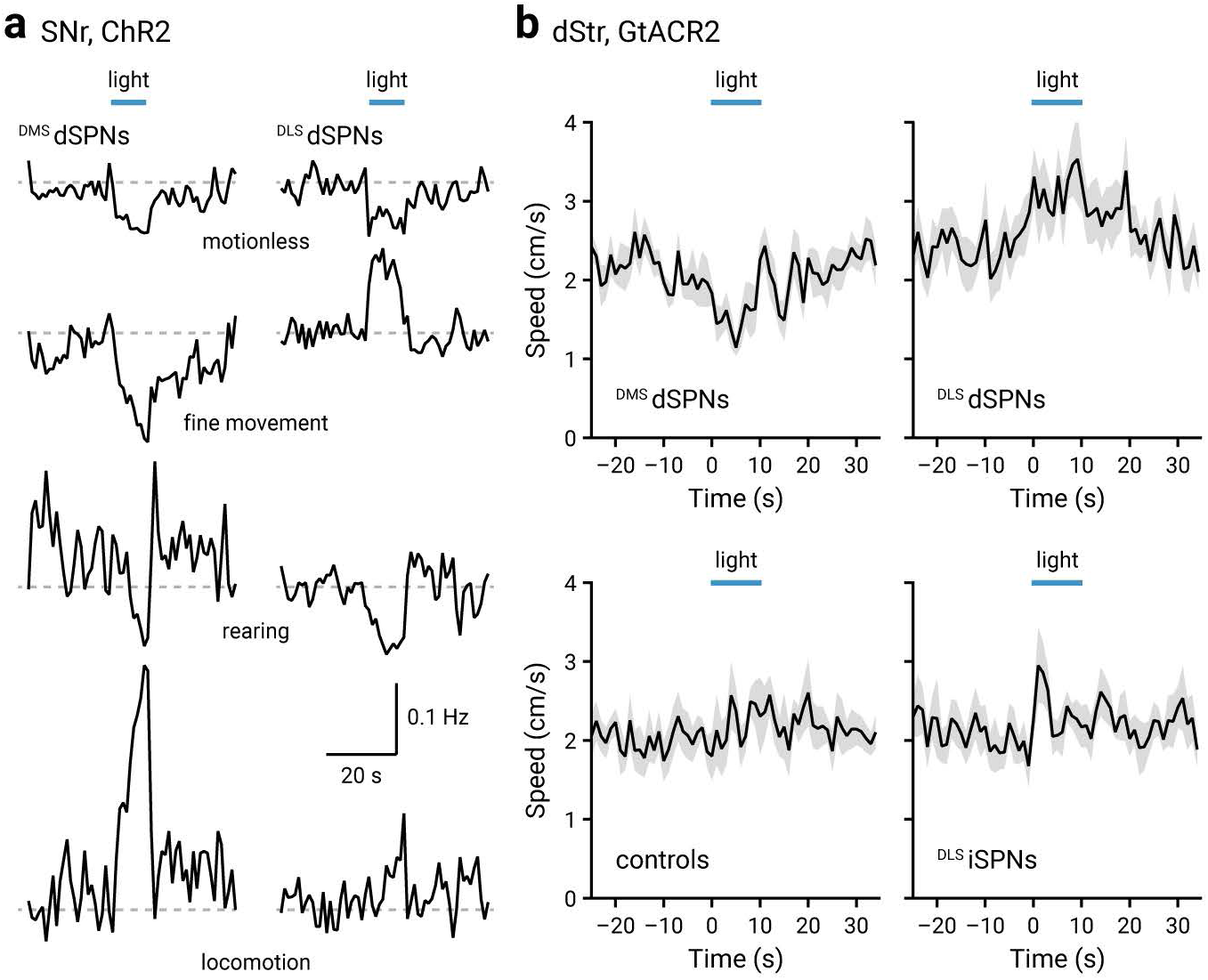
Extended optogenetic interrogation confirms distinct behavior roles of ^DMS^dSPNs and ^DLS^dSPNs. **a.** Mean changes in the event frequency of motionless, fine movement, rearing, and locomotion upon optogenetic stimulation (blue horizontal lines) of ^DMS^dSPNs and ^DLS^dSPNs terminals in the SNr. Scale bar applies to all traces. **b.** Changes in speeds are shown with light delivery (blue horizontal lines) in the DMS and DLS of *Drd1a*^Cre^ mice (top), DLS of Adora2a^Cre^ mice (bottom right) with GtACR2 expression. No changes in speeds was observed in the absence of opsin expression (bottom left).

The distinct behavioral patterns resulting from stimulation of the same SPN types from different spatial subdomains were surprising as it is commonly assumed that the role of the dStr in the regulation of locomotor activity is generalizable across DMS and DLS. To ensure that the inference made from our observations was not simply an artifact of gain-of-function experiments with ChR2, we performed additional optogenetic interrogation with an inhibitory opsin (i.e., GtACR2) (Mahn et al., 2018; Pamukcu et al., 2020). As expected from the diametric responses induced with optogenetic stimulation of ^DMS^dSPNs and ^DLS^dSPNs, GtACR2 activation in ^DMS^dSPNs and ^DLS^dSPNs induced suppression and promotion of speed, respectively (^DMS^dSPNs = –0.36 ± 0.13 fold, *n* = 8 mice, *P* = 0.016; ^DLS^dSPNs = 0.27 ± 0.16 fold, *n* = 11 mice, *P* = 0.0020) (**Figure 3b**). These responses were different (^DMS^dSPNs, *P* = 0.0037; ^DLS^dSPNs, *P* = 0.035) from those observed in mice with only light delivery (with no opsin expression to the DMS or DLS), which did not induce any consistent motor effects (0.04 ± 0.05 fold, *n* = 7 mice, *P* = 0.38). As optogenetic manipulations of ^DLS^dSPNs produced motor responses distinct from those of ^DMS^dSPNs, motor effects of optogenetic inhibition on ^DLS^iSPNs were examined. GtACR2 activation in ^DLS^iSPNs led to an increase in speed (^DLS^iSPNs = 0.15 ± 0.07 fold, *n* = 8 mice, *P* = 0.039), which contrasted with the motor promotion observed in ^DLS^dSPNs upon optogenetic inhibition.

### dSPNs send terminating axons to the GPe

We hypothesized that the observed behavioral responses were in part controlled at the GPe level where axons from both dSPNs and iSPNs converge (Chang et al., 1981; Kawaguchi et al., 1990; Wu et al., 2000; Levesque and Parent, 2005; Fujiyama et al., 2011; Cazorla et al., 2014; Okamoto et al., 2020). To appreciate the importance of the striatal input to the GPe relative to inputs from the rest of the brain, total synaptic inputs to the GPe were mapped. Two retrograde tracers, namely a Cre-expressing lentivirus (LVretro-Cre) (Knowland et al., 2017; Abecassis et al., 2020) and CTb 488 (Conte et al., 2009a, b) were co-injected into the GPe of a Cre-reporter (*R26*^LSL-tdTomato^) mouse. As expected, tdTomato^+^ and CTb 488^+^ neurons were readily observed in the dStr (**Figure 4a**), cross-validating the utility of the retrograde labeling strategy. Using serial two-photon tomography, whole-brain inputs to the GPe were mapped (Kim et al., 2017; Abecassis et al., 2020). As shown in **Figure 4a**, the dStr provides the largest input to the GPe. The number of input neurons (i.e., tdTomato^+^) from the dStr was at least an order of magnitude larger than that from other brain regions charted (e.g., central amygdala), constituting ∼80% (79.0 ± 3.1%) of total neurons (*n* = 45,223 neurons, 8 mice) that projected to the GPe. This observation is consistent with the earlier finding that GABAergic synapses amount to over 80% of all synapses in the GPe (Kita, 2007).

**Figure 4.**
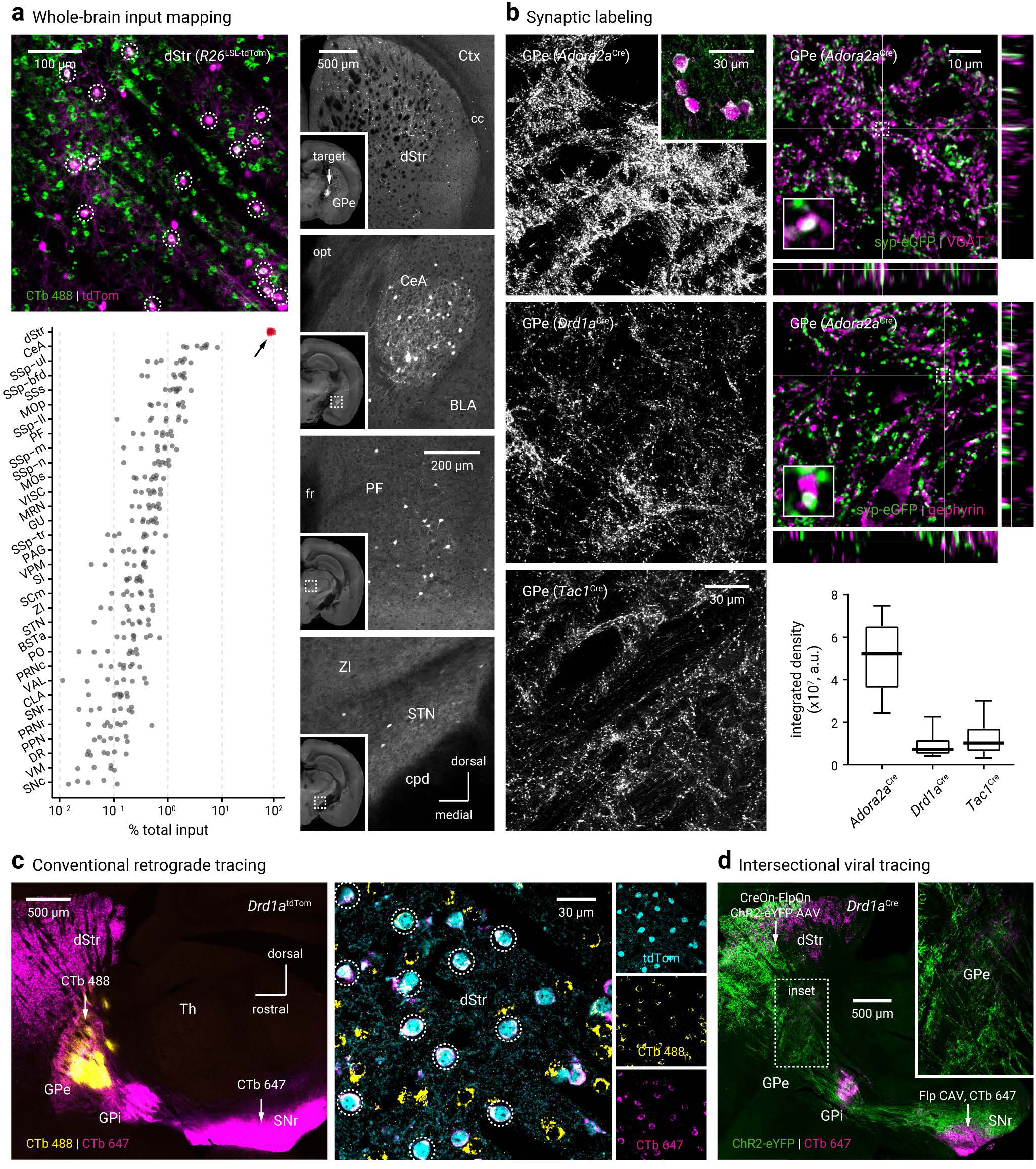
dSPNs send terminating axons to the GPe. **a.** *Top left*, a confocal micrograph showing retrogradely-labeled SPNs in the dStr from a Cre-reporter (*R26*^LSL-tdTomato^) mouse. A Cre-expressing lentivirus and CTb 488 were injected into the GPe. tdTomato^+^ and CTb 488^+^ neurons (in white circles) were visible. *Bottom left*, unbiased quantification of GPe-projecting neurons across the entire brain. Each marker represents a mouse (*n* = 8 mice). The arrow points to the data (red) from the dStr. *Right*, representative two-photon images from coronal sections showing GPe-projecting neurons were found in the dStr, central amygdala (CeA), parafascicular nucleus (PF), and subthalamic nucleus (STN). Inset in the first image indicates the location of the injection site. Scale bar in the third image applies to the bottom three images. **b.** *Left*, high-magnification images showing that terminals (eGFP^+^, white) from iSPNs or dSPNs were abundant in the GPe. CreOn-mRuby2-T2A-Synaptophysin-eGFP adeno-associated virus (AAV) was injected into the dStr of *Adora2a*^Cre^ (*top*), *Drd1a*^Cre^ (*middle*), and *Tac1*^Cre^ (*bottom*) mice to visualize SPN terminals. Maximal projections from ten optical sections are shown. *Inset*, mRuby2^+^ dSPNs in the dStr. *Top & middle right*, immunohistological analyses showing that eGFP^+^ boutons (green) were in proximity with vesicular GABA transporter (VGAT) and gephyrin (magenta), as shown in white in three orthogonal planes. Insets show magnified views of areas within the dotted square outlines. *Bottom right*, quantification of eGFP^+^ bouton density in the GPe from *Adora2a*^Cre^ (*n* = 12 ROIs), *Drd1a*^Cre^ (*n* = 10 ROIs), and *Tac1*^Cre^ (*n* = 12 ROIs) mice. **c.** *Left*, a low-magnification micrograph from a sagittal brain section demonstrating the target of retrograde tracers CTb 488 and CTb 647 into the GPe and SNr, respectively. The *Drd1a*^tdTomato^ allele was used to decipher the identity of SPNs. *Right*, representative high-magnification images illustrating CTb 488 and CTb 647 were detected in the same tdTom^+^ neurons in the dStr. White circles denote colocalization. **d.** A low-magnification micrograph from a sagittal brain section showing the expression of eYFP in the dStr as well as its projection targets including the GPe, GPi, and SNr following injections of CreOn-Flp canine adenovirus (CAV) into the SNr in combination with CreOn-FlpOn-ChR2-eYFP AAV into the dStr of a *Drd1a*^Cre^ mouse. CTb 647 was co-injected with CreOn-Flp CAV to visualize the injection site. Inset shows the eYFP^+^ axons in the GPe. Abbreviations: BLA, basolateral amygdalar nucleus; BSTa, bed nuclei of the stria terminalis, anterior division; cc, corpus callosum; CeA, central amygdala; CLA, claustrum; cpd, cerebral peduncle; Ctx, cortex; DR, dorsal raphe nucleus; dStr, dorsal striatum; fr, fasciculus retroflexus; GPe, external globus pallidus; GPi, internal globus pallidus; GU, gustatory areas; MOp, primary motor area; MOs, secondary motor area; MRN, midbrain reticular nucleus; opt, optic tract; PAG, periaqueductal gray; PF, parafascicular nucleus; PO, posterior complex of the thalamus; PPN, pedunculopontine nucleus; PRNc, pontine reticular nucleus, caudal part; PRNr, pontine reticular nucleus, rostral part; SCm, superior colliculus, motor related; SI, substantia innominata; SNc, substantia nigra pars compacta; SNr, substantia nigra pars reticulata; SSp-bfd, primary somatosensory area, barrel field; SSp-ll, primary somatosensory area, lower limb; SSp-m, primary somatosensory area, mouth; SSp-n, primary somatosensory area, nose; SSp-tr, primary somatosensory area, trunk; SSp-ul, primary somatosensory area, upper limb; SSs, supplemental somatosensory area; STN, subthalamic nucleus; VAL, ventral anterior-lateral complex of the thalamus; VISC, visceral area; VM, ventromedial thalamic nucleus; VPM, ventral posteromedial thalamic nucleus; Th, thalamus; ZI, zona incerta.

The whole-brain mapping approach did not take into account the number of synapses formed by each neuron type. To determine the innervation density of SPN subtypes, we injected into the dStr of *Adora2a*^Cre^ and *Drd1a*^Cre^ mice a Cre-inducible mRuby2-T2A-Synaptophysin-eGFP AAV, which tags transduced neurons and their axonal terminals with mRuby2 and eGFP, respectively (Knowland et al., 2017; Faget et al., 2018). As expected, eGFP^+^ puncta produced by SPNs were abundant in the GPe (**Figure 4b**). These eGFP^+^ puncta corresponded to GABAergic terminals, as demonstrated by their immunoreactivity for vesicular GABA transporter (VGAT) and gephyrin (**Figure 4b**). The abundance of eGFP^+^ puncta was thus used as a measure of innervation density. The density of eGFP^+^ puncta formed by iSPNs was about seven-fold higher than that formed by dSPNs (*Adora2a*^Cre^ = 5.2 ± 1.5 × 10^7^ a.u., *n* = 12 ROIs; *Drd1a*^Cre^ = 0.7 ± 0.3 × 10^7^ a.u., *n* = 10 ROIs; *P* < 0.0001) (**Figure 4b**). The disparity in the number of synaptic puncta formed by the two striatopallidal inputs is larger than that in earlier reports, which show that dSPNs provide roughly half the number of boutons compared to iSPNs in the GPe (Kawaguchi et al., 1990; Fujiyama et al., 2011).

To demonstrate whether single dSPNs innervate both the GPe and SNr, two different retrograde tracers (i.e., CTb 488 and CTb 647) were injected into the GPe and SNr, respectively. Using *Drd1a*^tdTomato^ mice to identify all dSPNs (Ade et al., 2011), we found that CTb 488 and CTb 647 signals were detected in the same tdTomato^+^ neurons within the dStr (**Figure 4c**). Using an intersectional approach (with *Drd1a*^Cre^ mice and a Flp-expressing retrograde virus) to confer an unparalleled spatial and genetic specificity, we observed axonal collateralization in the GPe from SNr-projecting, *Drd1a*^Cre+^ dStr neurons (**Figure 4d**). These findings confirmed the earlier observations from single-cell tracing studies that show SNr-projecting dStr neurons (i.e., dSPNs) arborize within the GPe (Kawaguchi et al., 1990; Wu et al., 2000; Levesque and Parent, 2005; Fujiyama et al., 2011). The results obtained from *Drd1a*^Cre^ and *Drd1a*^tdTomato^ mice were consistent with each other. Moreover, using the same synapse-tagging approach and CTb-based tracing mentioned above, we corroborated these findings in *Tac1*^Cre^ mice (eGFP^+^ puncta density = 1.0 ± 0.5 × 10^7^ a.u., *n* = 12 ROIs).

Though both *Drd1a*^Cre^ and *Adora2a*^Cre^ mice are well-characterized (Gong et al., 2007; Lobo et al., 2010; Gerfen et al., 2013; Okamoto et al., 2020), to ascertain that the observed striatopallidal innervation patterns were not simply artifacts from non-specific Cre recombination expression, we examined the arborization patterns and marker expression of the axons produced by Cre-expressing dStr neurons—they both gave the expected results and thus confirmed the validity of these two transgenic lines (**Figure 5a, b, & e**). Moreover, *ex vivo* recordings showed that dopamine D2 receptors selectively regulate striatopallidal GABA release from iSPN (*Adora2a*^Cre^) but not dSPN (*Drd1a*^Cre^) input (**Figure 5c & d**).

**Figure 5.**
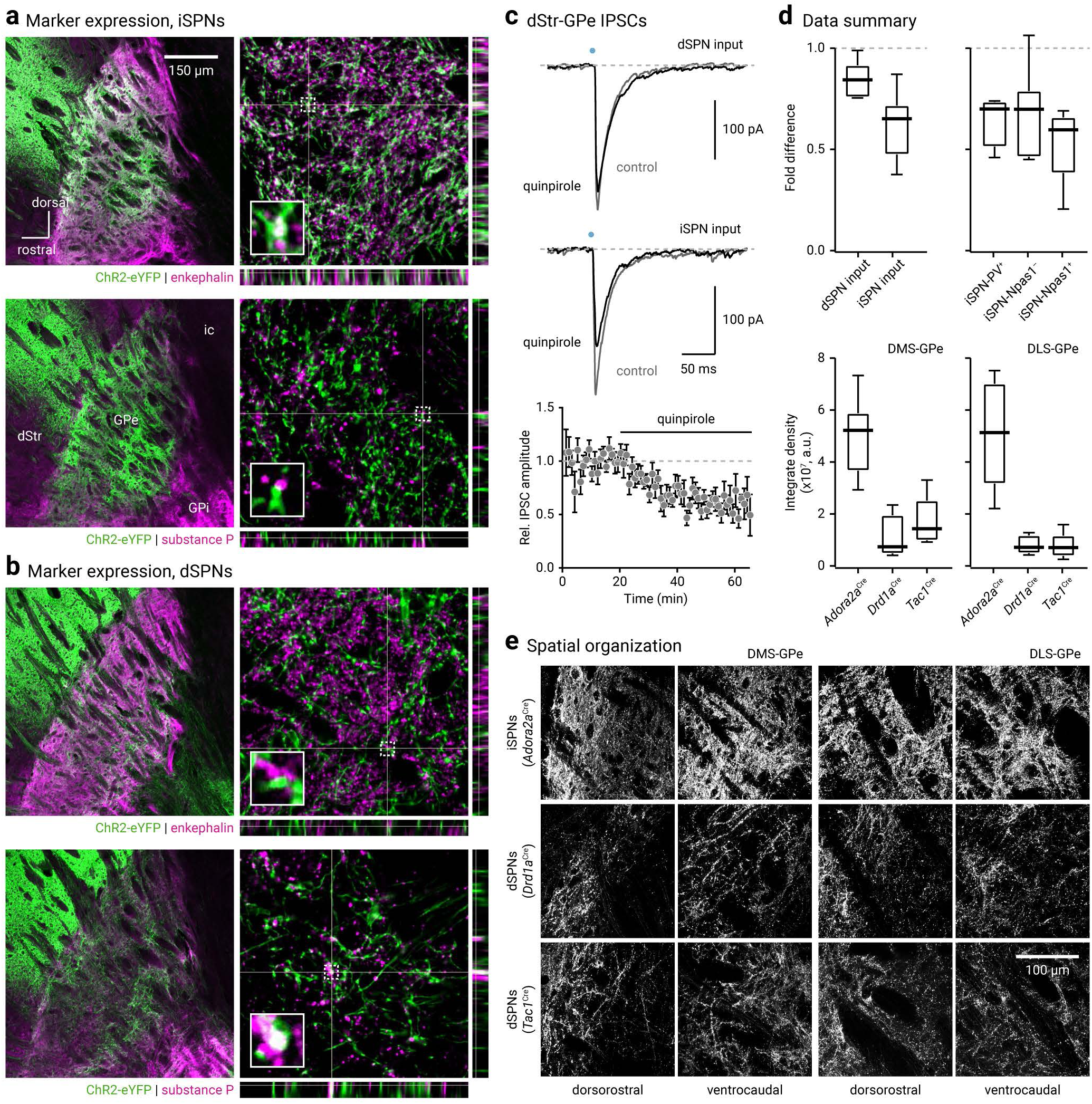
dSPN and iSPN inputs to GPe have unique properties. **a.** *Left*, confocal micrographs of two neighboring brain sections showing the iSPN projection to the GPe. The dStr of an *Adora2a*^Cre^ mouse was transduced with a CreOn-ChR2-eYFP AAV. The association of enkephalin (*top*) and substance P (*bottom*) with eYFP-labeled axonal fibers in the GPe was assessed with immunofluorescence labeling. *Right*, high-magnification images showing the spatial relationship between enkephalin (magenta, *top*) and substance P (magenta, *bottom*) with iSPN axons (green) in the GPe. Rectangular images show orthogonal projections. Crosshairs indicate the projected planes. Insets show magnified views of areas within the dotted square outlines. **b.** *Left*, confocal micrographs of two neighboring brain sections showing the dSPN projection to the GPe. The dStr of a *Drd1a*^Cre^ mouse was transduced with a CreOn-ChR2-eYFP AAV. The association of enkephalin (*top*) and substance P (*bottom*) with eYFP-labeled axonal fibers in the GPe was assessed with immunofluorescence labeling. *Right*, high-magnification images showing the spatial relationship between enkephalin (magenta, *top*) or substance P (magenta, *bottom*) and dSPN axons (green). Insets show magnified views of areas within the dotted square outlines. **c.** *Top*, two representative voltage-clamp recordings showing striatopallidal (dStr-GPe) inhibitory postsynaptic currents (IPSCs) in control (gray) condition and in the presence of quinpirole (10 µM, black). IPSCs were evoked with optogenetics; light was delivered in the dStr. *Drd1a*^Cre^ and *Adora2a*^Cre^ mice were used to examine the properties of dSPN and iSPN inputs, respectively. dSPN-GPe IPSCs (*top*) and iSPN-GPe IPSCs (*bottom*) are shown. The recordings were obtained from an Npas1^+^ and a PV^+^ neuron, respectively. *Bottom*, the relative iSPN-GPe IPSC amplitude was plotted vs. time (*n* = 12 neurons). The horizontal black line denotes the timing of quinpirole application. **d.** *Top*, box plots summarizing the effect of quinpirole on dStr-GPe IPSC amplitude. All recorded neurons for dSPN or iSPN input (dSPN input = 6 neurons, iSPN input = 16 neurons) are shown on the *left*. Plots on the *right* show iSPN input broken down by neuron types (PV^+^ = 4 neurons, Npas1^−^ = 7 neurons, Npas1^+^ = 5 neurons). *Bottom*, quantification of eGFP^+^ bouton density in the GPe shown in **e** (DMS: *Adora2a*^Cre^ = 6 ROIs, *Drd1a*^Cre^ = 4 ROIs, *Tac1*^Cre^ = 6 ROIs, DLS: *Adora2a*^Cre^ = 6 ROIs, *Drd1a*^Cre^ = 6 ROIs, *Tac1*^Cre^ = 6 ROIs). **e.** Representative high-magnification images showing SPN terminals (eGFP^+^, white) in the GPe. CreOn-mRuby2-T2A-Synaptophysin-eGFP AAV was injected into the dStr subregions of *Adora2a*^Cre^, *Drd1a*^Cre^ and *Tac1*^Cre^ mice. Terminals from iSPNs (*Adora2a*^Cre^) or dSPNs (*Drd1a*^Cre^ & *Tac1*^Cre^) from the DMS (*left*) or DLS (*right*) were abundant in the GPe. Images from medial and intermediate levels of the GPe are shown for terminals from the DMS and DLS, respectively. Maximal intensity from ten optical sections is shown for each example.

### Parallel organization of dStr-GPe-dStr loops

To understand if the organization of the striatopallidal projection formed the basis for the distinct motor outcomes associated with optogenetic stimulation of spatial subdomains within the dStr, we examined the anatomical organization of the striatal projections from the DMS and DLS. To study the dSPN and iSPN projections to the GPe, axonal terminals from dSPNs or iSPNs were selectively tagged using the synapse tagging approach (with synaptophysin-eGFP) described above. As shown in **Figure 5e**, the projections from the DMS and DLS were organized similarly at the GPe level; no systematic differences were observed along both dorsoventral and rostrocaudal axes. The innervation density of iSPN projection from the DMS and DLS, as measured by the abundance of eGFP^+^ puncta, were statistically indistinguishable (*Adora2a*^Cre^: DMS = 5.2 ± 0.7 × 10^7^ a.u., *n* = 6 ROIs; DLS = 5.1 ± 1.7 × 10^7^ a.u., *n* = 6 ROIs; *P* = 0.94) (**Figure 5d & e**). The dSPN projection estimated with *Drd1a*^Cre^ was organized similarly (*Drd1a*^Cre^: DMS = 0.7 ± 0.2 × 10^7^ a.u., *n* = 4 ROIs; DLS = 0.7 ± 0.2 × 10^7^ a.u., *n* = 6 ROIs; *P* = 0.91) (**Figure 5d & e**). Using *Tac1*^Cre^ to estimate the dSPN projection to the GPe, we found a stronger projection from the DMS than DLS (*Tac1*^Cre^: DMS = 1.4 ± 0.5 × 10^7^ a.u., *n* = 6 ROIs; DLS = 0.7 ± 0.3 × 10^7^ a.u., *n* = 6 ROIs; *P* = 0.041) (**Figure 5d & e**). We do not currently have an explanation for the difference observed between *Drd1a*^Cre^ and *Tac1*^Cre^ mice; it is possible that these two lines have nuanced differences in the striosome-matrix bias.

On the contrary, axons of dSPNs and iSPNs from the DMS and DLS target the medial and lateral portions of their downstream targets, respectively. Moreover, the projections from iSPNs and dSPNs appeared to follow the same topographical pattern (**Figure 6a & b**). These results were consistent with the functional anatomy dissected using optogenetic approaches (**Figure 1h**). In sum, our findings contextualize earlier observations that striatofugal axons are topographically arranged with new cellular insights (Szabo, 1962; Cowan and Powell, 1966; Chang et al., 1981; Wilson and Phelan, 1982; Hedreen and DeLong, 1991; Deniau et al., 1996; Romanelli et al., 2005; Fujiyama et al., 2011; Nambu, 2011; Bertino et al., 2020; Foster et al., 2020; Lee et al., 2020; Okamoto et al., 2020). As SPN axons did not appear to run extensively along the mediolateral axis (**Figure 6a & b**), we asked if SPNs from distinct spatial domains could interact through the feedback projections from the GPe (Abdi et al., 2015; Dodson et al., 2015; Hernandez et al., 2015; Glajch et al., 2016; Hegeman et al., 2016; Mallet et al., 2016; Abecassis et al., 2020). To this end, we labeled striatally projecting GPe neurons retrogradely using CTb 647. As shown in **Figure 6c**, DMS- and DLS-projecting neurons resided in distinct spatial domains within the GPe, demonstrating a parallel organization of dStr-GPe-dStr loops.

**Figure 6.**
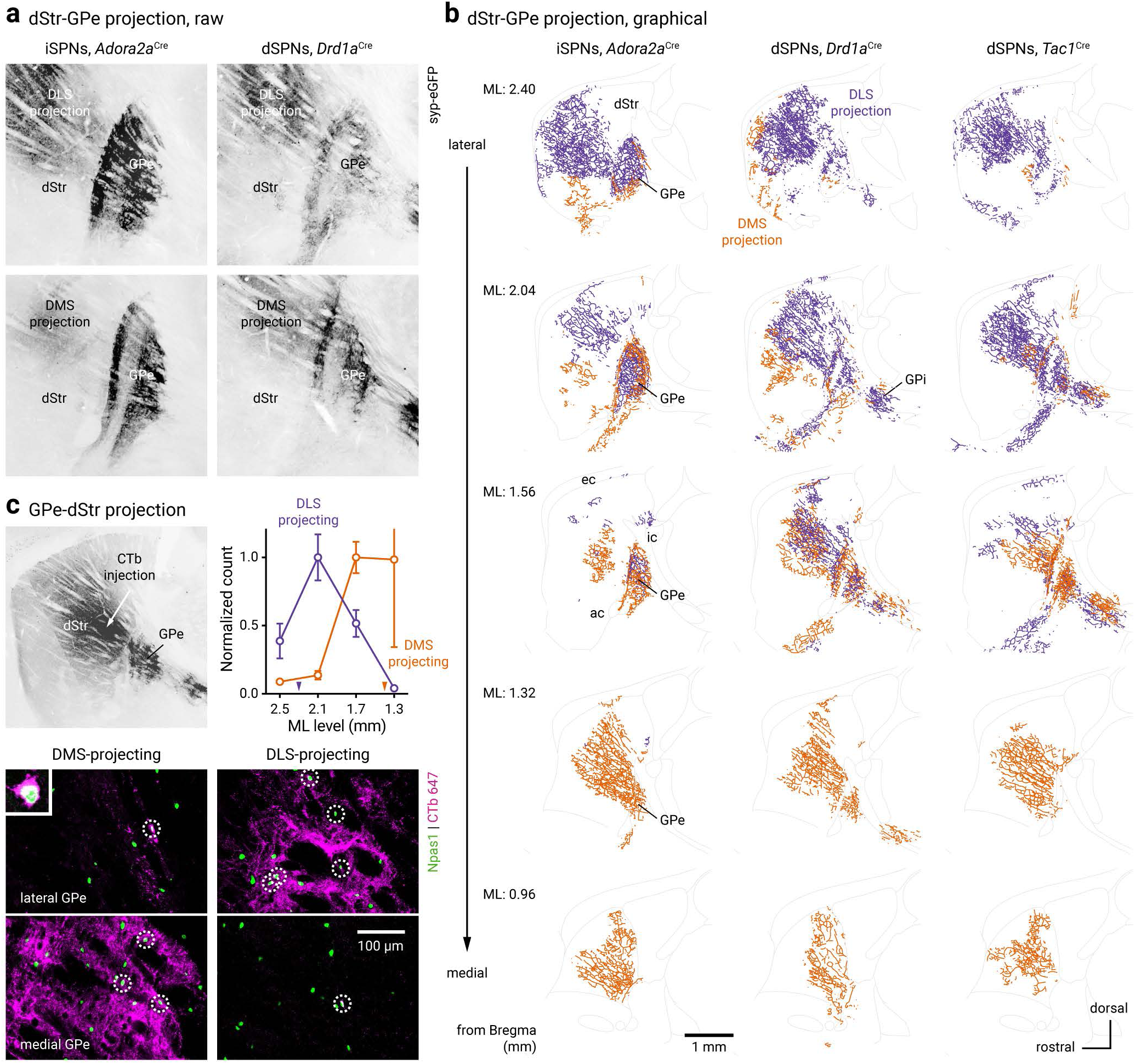
Topographical organization of dStr-GPe-dStr projections. **a.** Representative epifluorescence images showing eGFP signals from CreOn-mRuby2-T2A-Synaptophysin-eGFP AAV injections into the DMS or DLS of *Adora2a*^Cre^ (left) and *Drd1a*^Cre^ (right) mice. **b.** Graphical representation of terminals from iSPNs and dSPNs across five different sagittal planes. AAV was injected into *Adora2a*^Cre^ (left), *Drd1a*^Cre^ (middle), and *Tac1*^Cre^ (right) mice. Low-magnification epifluorescence images (see **a**) of eGFP-labeled terminals were vectorized. Terminals from the DMS and DLS are represented by orange and purple line segments, respectively. Scale bar applies to all panels. Abbreviations: ac, anterior commissure; ec, external capsule, GPi, internal globus pallidus; ic, internal capsule. **c.** *Top left*, a low magnification image showing fluorescence signals following a striatal CTb 647 injection. *Top right*, Normalized counts (mean ± sem) for CTb-labelled (CTb^+^) Npas1^+^ neurons in the GPe are shown. Neurons were retrogradely-labelled from DMS (orange, *n* = 148 neurons, 12 sections) and DLS (purple, *n* = 241 neurons, 12 sections) injections. Injection locations are marked by the arrowheads on the horizontal-axis. *Bottom*, CTb^+^ (magenta) and Npas1^+^ (green) neurons were seen in the GPe. White circles denote co-labeling. Inset, a representative example showing the co-labeling of CTb 647 and Npas1 in a GPe neuron.

### ^DLS^dSPNs strongly target Npas1^+^ neurons

To determine the postsynaptic targets of SPN subtypes in the GPe, we used rabies virus (RbV)-mediated tracing (Hunt et al., 2018; Shin et al., 2018). Among all SPNs that were retrogradely labeled from PV^+^ neurons, a larger fraction were positive for *Drd2* mRNA than for *Drd1a* mRNA (*Drd1a* = 40.7 ± 3.1%, *Drd2* = 59.3 ± 3.1%, *n* = 5 mice) (**Figure 7a & c**). Conversely, among all SPNs that were retrogradely labeled from Npas1^+^ neurons, two-thirds were positive for *Drd1a* mRNA (*Drd1a* = 66.4 ± 0.6%, *Drd2* = 33.6 ± 0.6%, *n* = 5 mice) (**Figure 7a & c**); this connectivity pattern was different from the relative abundance of *Drd1a-* or *Drd2*-expressing SPNs that project to the PV^+^ neurons (*P* < 0.0001).

**Figure 7.**
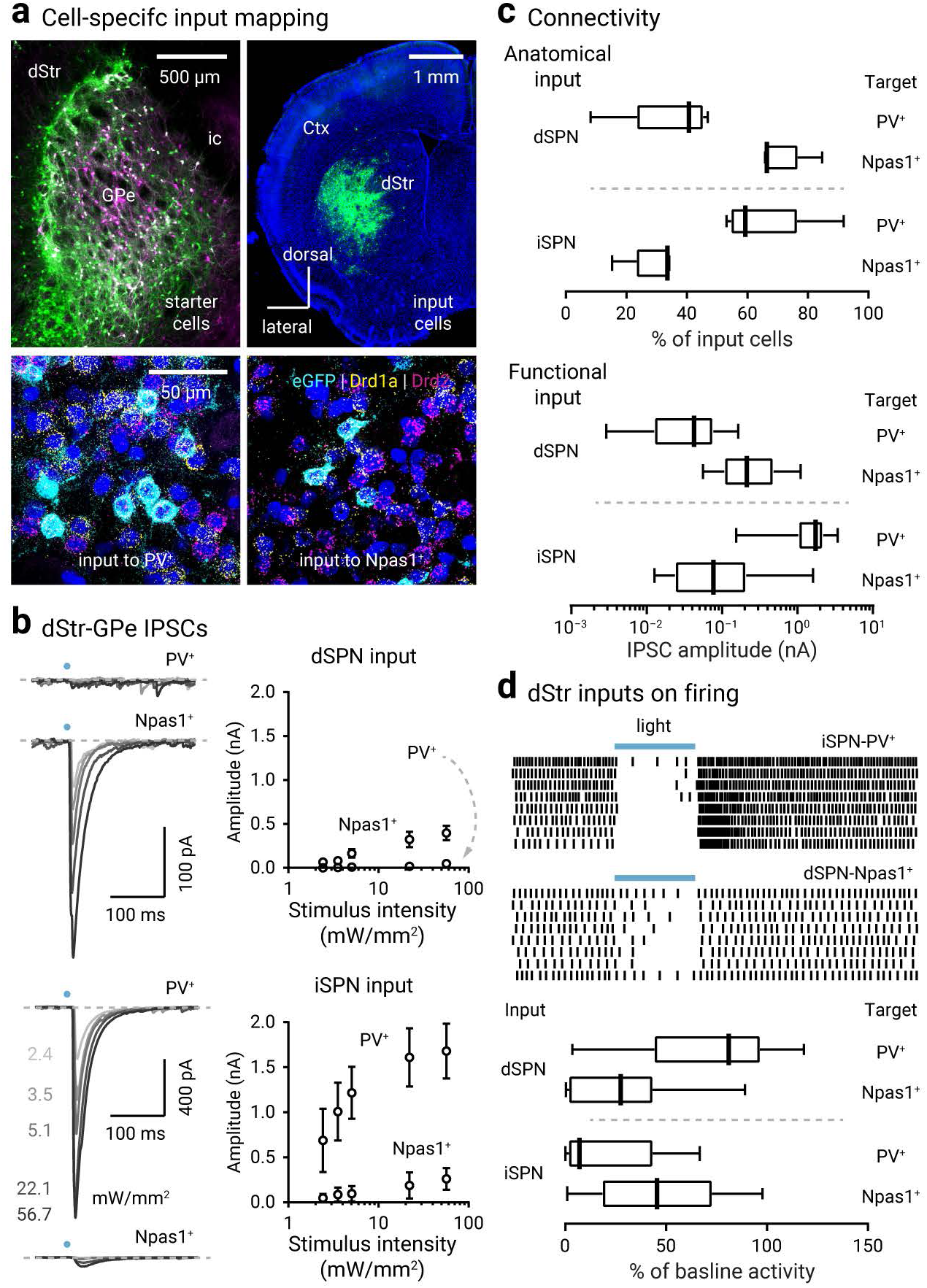
dSPNs strongly innervate Npas1^+^ neurons. **a.** *Top*, low-magnification images showing GPe starter cells (*left,* white) and striatal input cells (*right,* green) in coronal brain sections from *Npas1*^Cre^ (*left*) or *Pvalb*^Cre^ (*right*) mice injected with hSyn-DIO-mRuby2-TVA-RVG AAV and rabies virus expressing eGFP sequentially into the GPe. *Bottom*, representative high-magnification images showing that striatal input cells (eGFP^+^, green) from *Pvalb*^Cre^ (*left*) or *Npas1*^Cre^ (*right*) mice were positive for *Drd1a* (yellow) or *Drd2* (magenta) mRNA. **b.** *Left*, representative traces showing IPSCs recorded in PV^+^ or Npas1^+^ neurons with optogenetic stimulation of dSPNs (*top*) or iSPNs (*bottom*) in the DLS. Traces from five stimulus intensities (2.4–56.7 mW/mm^2^) are shown. *Right*, input-output relationship for corresponding inputs. A full list of median values, sample sizes, and statistical comparisons at different stimulus intensities for discrete inputs is shown in **Table 3**. Response rate for each input is shown in **Table 5**. **c.** Summary of anatomical (see **a**; *Pvalb*^Cre^ = 5 mice, *Npas1*^Cre^ = 5 mice) and functional (see **b**; dSPN-PV^+^ = 8 neurons, dSPN-Npas1^+^ = 28 neurons, iSPN-PV^+^ = 12 neurons, iSPN-Npas1^+^ = 20 neurons) connectivity for discrete dStr-GPe inputs. The IPSC amplitudes at maximal stimulus intensity are plotted. **d.** *Top*, representative raster plots showing that stimulation (10 Hz for 2 s) of iSPNs or dSPNs in the DLS strongly suppressed firing of PV^+^ or Npas1^+^ neurons, respectively. Blue bars indicate the period of blue light stimulation. *Bottom*, summary of changes in baseline activity with stimulation of discrete dStr-GPe inputs (dSPN-PV^+^ = 9 neurons, dSPN-Npas1^+^ = 11 neurons, iSPN-PV^+^ = 9 neurons, iSPN-Npas1^+^ = 11 neurons).

**Table 3.** Input-output relationship of DLS inputs in naïve and 6-OHDA lesioned mice.

| Input & stimulus location | Stimulus intensity (mW/mm<sup>2</sup>) | IPSC peak amplitude (pA) |  |  |  |  |  |  |  | Statistical analyses <sup>b</sup> |  |  |  |
| --- | --- | --- | --- | --- | --- | --- | --- | --- | --- | --- | --- | --- | --- |
|  |  | naïve PV<sup>+</sup> |  | naïve Npas1<sup>+</sup> |  | 6-OHDA PV<sup>+</sup> |  | 6-OHDA Npas1<sup>+</sup> |  | naïve PV<sup>+</sup> vs naïve Npas1<sup>+</sup> | naïve PV<sup>+</sup> vs 6-OHDA PV<sup>+</sup> | naïve Npas1<sup>+</sup> vs 6-OHDA Npas1<sup>+</sup> | 6-OHDA PV<sup>+</sup> vs 6-OHDA Npas1<sup>+</sup> |
|  |  | Median ± MAD | *n*<sup>a</sup> | Median ± MAD | *n* | Median ± MAD | *n* | Median ± MAD | *n* |  |  |  |  |
| dSPN, dStr | 2.4 | 0.0 ± 0.0 | 4 | 22.7 ± 22.7 | 11 | 2.0 ± 2.0 | 10 | 324.2 ± 290.8 | 11 | 0.078 | 0.27 | **0.0052** | **0.0046** |
|  | 3.5 | 2.1 ± 2.1 | 5 | 33.7 ± 33.7 | 12 | 5.6 ± 5.6 | 12 | 271.1 ± 255.5 | 12 | 0.076 | 0.48 | **0.0044** | **0.0035** |
|  | 5.1 | 8.1 ± 6.2 | 6 | 79.1 ± 59.5 | 17 | 10.5 ± 10.5 | 13 | 375.5 ± 305.5 | 13 | **0.00014** | 0.62 | **0.017** | **0.0050** |
|  | 22.1 | 7.7 ± 3.5 | 8 | 158.3 ± 125.0 | 18 | 55.4 ± 54.1 | 13 | 866.3 ± 537.5 | 13 | **< 0.0001** | **0.037** | **0.0095** | **0.0051** |
|  | 56.7 | 38.5 ± 29.3 | 8 | 213.8 ± 122.0 | 28 | 81.0 ± 76.7 | 13 | 815.8 ± 530.5 | 15 | **0.00011** | 0.14 | **0.0017** | **0.0015** |
| dSPN, GPe | 2.1 | 15.7 ± 12.6 | 12 | 182.7 ± 170.1 | 20 | 120.0 ± 102.1 | 11 | 527.7 ± 447.2 | 23 | **0.012** | 0.051 | **0.039** | **0.042** |
|  | 3.5 | 43.9 ± 36.3 | 12 | 448.4 ± 371.0 | 20 | 364.7 ± 310.1 | 11 | 1,208.7 ± 818.2 | 23 | **0.0031** | **0.044** | **0.013** | **0.0075** |
| iSPN, dStr | 2.4 | 295.0 ± 253.8 | 6 | 0.0 ± 0.0 | 13 | 578.9 ± 503.9 | 10 | 4.3 ± 4.3 | 6 | **0.00085** | 0.79 | 0.34 | **0.023** |
|  | 3.5 | 761.7 ± 439.2 | 8 | 0.0 ± 0.0 | 14 | 819.5 ± 667.0 | 13 | 17.1 ± 17.1 | 7 | **0.00041** | 0.86 | 0.32 | **0.0011** |
|  | 5.1 | 969.2 ± 546.4 | 10 | 8.6 ± 7.7 | 15 | 891.6 ± 544.2 | 14 | 12.7 ± 5.6 | 8 | **0.00022** | 0.71 | 0.40 | **0.00028** |
|  | 22.1 | 1,593.1 ± 682.9 | 11 | 30.0 ± 23.1 | 14 | 1,355.1 ± 713.5 | 16 | 29.8 ± 19.2 | 10 | **0.00052** | 0.75 | 0.80 | **< 0.0001** |
|  | 56.7 | 1,740.7 ± 627.7 | 12 | 76.9 ± 60.9 | 20 | 1,682.0 ± 820.4 | 18 | 39.2 ± 22.2 | 11 | **< 0.0001** | 0.75 | 0.36 | **< 0.0001** |
| iSPN, GPe | 2.1 | 1,495.6 ± 440.9 | 16 | 103.4 ± 74.6 | 16 | 1,705.2 ± 480.2 | 16 | 57.8 ± 43.9 | 15 | **0.0034** | 0.24 | 0.13 | **< 0.0001** |
|  | 3.5 | 1,715.2 ± 691.9 | 16 | 288.0 ± 212.9 | 17 | 1,806.1 ± 689.2 | 16 | 94.5 ± 84.0 | 15 | **0.0016** | 0.36 | 0.19 | **< 0.0001** |

Notes:
<sup>a</sup> Sample size (*n*) only included cells that responded to maximal stimulus intensity.<sup>b</sup> Mann-Whitney *U* test; two-tailed exact *P* values were shown. *P* < 0.05 were shown in bold.

The anatomical data showed a biased connectivity pattern where PV^+^ neurons and Npas1^+^ neurons were preferentially innervated by iSPNs and dSPNs, respectively. However, we were precluded from interrogating the spatial organization of the striatopallidal subcircuits with RbV-tracing, as the GPe is relatively small along the mediolateral axis. Instead, we performed *ex vivo* patch-clamp recordings in identified GPe neurons. The size of optogenetically evoked inhibitory postsynaptic currents (IPSCs) was used as a measure of connection strength. The strength of ^DLS^dSPN input to PV^+^ neurons was smaller than that to Npas1^+^ neurons. This difference was consistently observed across a large range of stimulus intensities (5.1–56.7 mW/mm^2^) (IPSC_max_: PV^+^ = 38.5 ± 29.3 pA, *n* = 8 neurons; Npas1^+^ = 213.8 ± 122.0 pA, *n* = 28 neurons; *P* = 0.00011) (**Figure 7b & c**). These electrophysiological and anatomical data collectively argue that ^DLS^dSPNs targeted Npas1^+^ neurons over PV^+^ neurons.

Our RbV-tracing data suggest that iSPNs preferentially innervate PV^+^ neurons. Here we asked whether ^DLS^iSPNs differentially target PV^+^ neurons versus Npas1^+^ neurons. As shown in **Figure 7b & c**, PV^+^ neurons received stronger iSPN input compared to Npas1^+^ neurons; this difference was observed across a wide range of stimulus intensities (2.4–56.7 mW/mm^2^) (IPSCs_max_: PV^+^ = 1,740.7 ± 627.7 pA, *n* = 12 neurons; Npas1^+^ = 76.9 ± 60.9 pA, *n* = 20 neurons; *P* < 0.0001). Among all individual DLS inputs examined, ^DLS^iSPN-PV^+^ input was the strongest. In summary, our data indicate that ^DLS^iSPNs preferentially target PV^+^ neurons. A full description of the input-output relationship for distinct DLS inputs is summarized in **Table 3**.

Consistent with the differences in ^DLS^dSPN input between PV^+^ neurons and Npas1^+^ neurons, optogenetic stimulation of ^DLS^dSPNs only modestly decreased the firing of PV^+^ neurons, but strongly suppressed the firing of Npas1^+^ neurons (PV^+^ = –0.19 ± 0.21 fold, *n* = 9 neurons, *P* = 0.027; Npas1^+^ = –0.73 ± 0.23 fold, *n* = 11 neurons, *P* = 0.00098) (**Figure 7d**); the effects were different between PV^+^ neurons and Npas1^+^ neurons (*P* = 0.020). On the other hand, although there was a big difference in the strength of the ^DLS^iSPN input between PV^+^ neurons and Npas1^+^ neurons, we did not observe a difference (*P* = 0.083) in the fold change of firing between PV^+^ neurons and Npas1^+^ neurons from ^DLS^iSPN stimulation (PV^+^ = –0.93 ± 0.06 fold, *n* = 9 neurons, *P* = 0.0039; Npas1^+^ = –0.55 ± 0.27 fold, *n* = 11 neurons, *P* = 0.0020) (**Figure 7d**). Given the higher input resistance of Npas1^+^ neurons (Hernandez et al., 2015), it is not surprising to see that a weak ^DLS^iSPN input to Npas1^+^ neurons suppressed firing disproportionally. The GABAergic nature of the ^DLS^dSPN input was confirmed in a subset of Npas1^+^ neurons. Similarly, in all PV^+^ neurons tested (*n* = 5 neurons), the application of SR95531 completely blocked the ^DLS^iSPN IPSCs, confirming the GABAergic nature of the synapse (**Figure 8c**).

**Figure 8.**
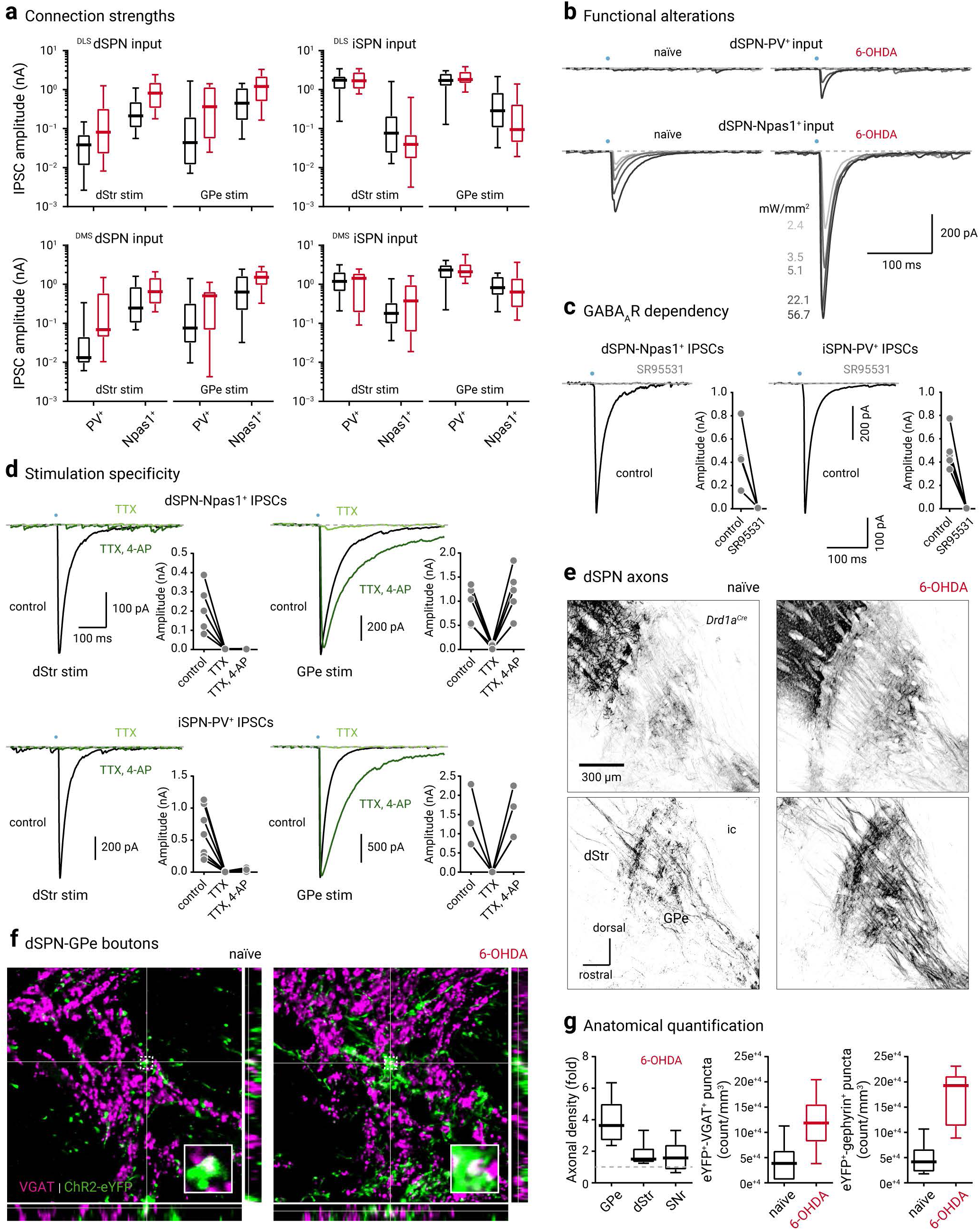
dSPN inputs are selectively strengthened following chronic 6-OHDA lesion. **a.** Connection strengths from dSPNs (*left*) and iSPNs (*right*) to PV^+^ neurons and Npas1^+^ neurons across two DLS (*top*) and DMS domains (*bottom*) were assessed with both somatic (i.e., intrastriatal, dStr) and terminal (i.e., intrapallidal, GPe) stimulation. Data from naïve (black) and 6-OHDA lesioned (red) mice are shown. Results are summarized as box plots. See **Table 3 & 4** for a full listing of IPSC amplitudes and sample sizes. **b.** Representative voltage-clamp recordings showing IPSCs arose from dSPNs in PV^+^ neurons (*top*) and Npas1^+^ neurons (*bottom*). A range of stimulus intensities (2.4–56.7 mW/mm^2^) were tested. Note the increase in IPSC amplitude in 6-OHDA mice (red, *right*) compared to that from naïve (black, *left*). **c.** GABA_A_ receptor dependency of the IPSCs were tested with a GABA_A_ receptor antagonist, SR95531 (10 µM). Each marker represents a cell (dSPN-Npas1^+^ = 4 neurons, iSPN-PV^+^ = 5 neurons). **d.** The spatial specificity of the optogenetic stimulation was assessed with the applications of tetrodotoxin (TTX 1 µM, light green) and its co-application with 4-aminopyridine (4-AP 100 µM, dark green). Each marker represents a cell (dSPN-Npas1^+^: dStr stim = 5 neurons, GPe stim = 5 neurons, iSPN-PV^+^: dStr stim = 7 neurons, GPe stim = 3 neurons). **e.** Confocal micrographs showing the innervation of dSPN axons in the GPe from naïve (*left*) and 6-OHDA lesioned (*right*) mice. To visualize dSPN axons, *Drd1a*^Cre^ mice were transduced with CreOn-ChR2-eYFP AAV. Intermediate (*top*) and medial (*bottom*) levels are shown. **f.** Representative high-magnification images showing dSPN bouton density in the GPe from naïve (*left*) and 6-OHDA lesioned (*right*) mice. dSPN boutons were represented by VGAT^+^ (magenta) and ChR2-eYFP^+^ (green) puncta. Breakout panels show orthogonal xz-projection (*bottom*) and yz-projection (*right*). Crosshairs indicate the pixel of interest. The colocalization of the signals is shown as white. Insets show magnified views of areas within the dotted square outlines. **g.** Quantification of the data shown in **e** and **f**: axonal density (naïve = 4 mice, 6-OHDA = 6 mice), eYFP^+^-VGAT^+^ puncta density (naïve = 6 sections, 6-OHDA = 10 sections), and eYFP^+^-gephyrin^+^ puncta density (naïve = 6 sections, 6-OHDA = 8 sections).

**Table 4.** Input-output relationship of DMS inputs in naïve and 6-OHDA lesioned mice.

| Input & stimulus location | Stimulus intensity (mW/mm<sup>2</sup>) | IPSC peak amplitude (pA) |  |  |  |  |  |  |  | Statistical analyses <sup>b</sup> |  |  |  |
| --- | --- | --- | --- | --- | --- | --- | --- | --- | --- | --- | --- | --- | --- |
|  |  | naïve PV<sup>+</sup> |  | naïve Npas1<sup>+</sup> |  | 6-OHDA PV<sup>+</sup> |  | 6-OHDA Npas1<sup>+</sup> |  | naïve PV<sup>+</sup> vs naïve Npas1<sup>+</sup> | naïve PV<sup>+</sup> vs 6-OHDA PV<sup>+</sup> | naïve Npas1<sup>+</sup> vs 6-OHDA Npas1<sup>+</sup> | 6-OHDA PV<sup>+</sup> vs 6-OHDA Npas1<sup>+</sup> |
|  |  | Median ± MAD | *n*<sup>a</sup> | Median ± MAD | *n* | Median ± MAD | *n* | Median ± MAD | *n* |  |  |  |  |
| dSPN, dStr | 2.4 | 8.5 ± 8.5 | 8 | 16.2 ± 16.2 | 15 | 16.3 ± 16.3 | 14 | 109.5 ± 107.4 | 20 | 0.17 | 0.53 | 0.23 | **0.012** |
|  | 3.5 | 3.8 ± 3.8 | 7 | 17.3 ± 17.3 | 18 | 37.1 ± 32.8 | 14 | 88.6 ± 84.9 | 21 | 0.18 | 0.22 | 0.068 | **0.047** |
|  | 5.1 | 5.9 ± 4.6 | 12 | 48.7 ± 46.2 | 18 | 35.4 ± 28.7 | 15 | 144.3 ± 124.7 | 27 | **0.021** | **0.046** | 0.052 | **0.0019** |
|  | 22.1 | 11.2 ± 7.7 | 14 | 123.0 ± 91.2 | 22 | 87.7 ± 74.2 | 15 | 498.5 ± 289.9 | 27 | **0.00051** | **0.0079** | **0.0019** | **0.00053** |
|  | 56.7 | 13.3 ± 6.5 | 16 | 246.2 ± 156.7 | 23 | 69.1 ± 56.0 | 16 | 649.7 ± 396.5 | 27 | **< 0.0001** | **0.0085** | **0.0093** | **0.00044** |
| dSPN, GPe | 2.1 | 33.8 ± 28.2 | 17 | 469.5 ± 334.9 | 21 | 272.2 ± 261.7 | 19 | 1,138.4 ± 664.5 | 27 | **0.0013** | 0.17 | **0.030** | **< 0.0001** |
|  | 3.5 | 75.7 ± 64.7 | 17 | 635.6 ± 551.0 | 19 | 507.1 ± 384.7 | 17 | 1,518.0 ± 685.4 | 24 | **0.0048** | 0.25 | **0.033** | **< 0.0001** |
| iSPN, dStr | 2.4 | 14.5 ± 13.2 | 19 | 23.5 ± 23.5 | 19 | 31.3 ± 29.3 | 11 | 28.5 ± 28.3 | 16 | 0.74 | 0.66 | 0.68 | 0.58 |
|  | 3.5 | 11.9 ± 11.4 | 22 | 34.2 ± 31.8 | 20 | 236.5 ± 216.5 | 18 | 31.6 ± 31.6 | 19 | 0.76 | **0.011** | 0.88 | **0.031** |
|  | 5.1 | 43.4 ± 39.6 | 22 | 36.6 ± 35.2 | 20 | 282.0 ± 271.5 | 22 | 62.7 ± 42.9 | 17 | 0.42 | 0.16 | 0.56 | 0.11 |
|  | 22.1 | 674.9 ± 548.4 | 25 | 136.1 ± 95.6 | 23 | 1,009.3 ± 864.9 | 24 | 193.7 ± 172.0 | 19 | **0.016** | 0.41 | 0.35 | **0.038** |
|  | 56.7 | 1,191.7 ± 640.5 | 25 | 179.8 ± 89.5 | 24 | 1,433.9 ± 890.2 | 25 | 376.5 ± 339.4 | 20 | **< 0.0001** | 0.51 | 0.52 | **0.021** |
| iSPN, GPe | 2.1 | 2,185.5 ± 819.1 | 40 | 625.0 ± 459.1 | 24 | 1,889.3 ± 999.3 | 25 | 506.5 ± 410.5 | 19 | **< 0.0001** | 0.84 | 0.78 | **0.0011** |
|  | 3.5 | 2,340.6 ± 849.7 | 41 | 823.0 ± 442.4 | 25 | 2,098.5 ± 727.5 | 25 | 635.5 ± 417.9 | 17 | **< 0.0001** | 0.83 | 0.46 | **0.00025** |

Notes:
<sup>a</sup> Sample size (*n*) only included cells that responded to maximal stimulus intensity.<sup>b</sup> Mann-Whitney *U* test; two-tailed exact *P* values were shown. *P* < 0.05 were shown in bold.

**Table 5.** Response rate to discrete dStr inputs in naïve and 6-OHDA lesioned mice.

| Input | Stimulus location | Stimulus intensity (mW/mm<sup>2</sup>) | Percentage of responders (%) |  |  |  |  |  |  |  | Statistical analyses <sup>b</sup> |  |  |  |
| --- | --- | --- | --- | --- | --- | --- | --- | --- | --- | --- | --- | --- | --- | --- |
|  |  |  | naïve PV<sup>+</sup> |  | naïve Npas1<sup>+</sup> |  | 6-OHDA PV<sup>+</sup> |  | 6-OHDA Npas1<sup>+</sup> |  | naïve PV<sup>+</sup> vs naïve Npas1<sup>+</sup> | naïve PV<sup>+</sup> vs 6-OHDA PV<sup>+</sup> | naïve Npas1<sup>+</sup> vs 6-OHDA Npas1<sup>+</sup> | 6-OHDA PV<sup>+</sup> vs 6-OHDA Npas1<sup>+</sup> |
|  |  |  | % | *n*<sup>a</sup> | % | *n* | % | *n* | % | *n* |  |  |  |  |
| <sup>DLS</sup>dSPN | dStr | 56.7 | 44.4 | 18 | 93.8 | 32 | 86.7 | 15 | 100 | 15 | **0.00019** | **0.027** | > 0.99 | 0.48 |
|  | GPe | 3.5 | 70.6 | 17 | 95.2 | 21 | 91.7 | 12 | 100 | 18 | 0.071 | 0.35 | > 0.99 | 0.40 |
| <sup>DLS</sup>iSPN | dStr | 56.7 | 92.9 | 14 | 90.9 | 22 | 100 | 18 | 68.8 | 16 | > 0.99 | 0.44 | 0.11 | **0.016** |
|  | GPe | 3.5 | 100 | 9 | 94.4 | 18 | 100 | 16 | 100 | 15 | > 0.99 | > 0.99 | > 0.99 | > 0.99 |
| <sup>DMS</sup>dSPN | dStr | 56.7 | 76.2 | 21 | 100 | 24 | 61.5 | 26 | 100 | 30 | **0.017** | 0.35 | > 0.99 | **0.00015** |
|  | GPe | 3.5 | 94.4 | 18 | 100 | 21 | 95 | 20 | 100 | 28 | 0.46 | > 0.99 | > 0.99 | 0.42 |
| <sup>DMS</sup>iSPN | dStr | 56.7 | 100 | 25 | 89.7 | 29 | 100 | 26 | 96.2 | 26 | 0.24 | > 0.99 | 0.61 | > 0.99 |
|  | GPe | 3.5 | 100 | 41 | 100 | 25 | 100 | 25 | 100 | 19 | > 0.99 | > 0.99 | > 0.99 | > 0.99 |

Notes:
<sup>a</sup> Sample size (*n*) included all cells with or without responses.
<sup>b</sup> Fisher's exact test; two-tailed exact *P* values were shown. *P* < 0.05 were shown in bold.

### DMS and DLS inputs share a similar organization

So far, we have shown that ^DLS^dSPNs and ^DLS^iSPNs preferentially innervate Npas1^+^ neurons and PV^+^ neurons, respectively. While there are subtle spatial distributions of neuron subtypes in the GPe (Mastro et al., 2014; Abdi et al., 2015; Hernandez et al., 2015; Abecassis et al., 2020), their relationship with striatopallidal projections was not known. To provide insights into whether the organization of the striatopallidal projections across the mediolateral axis could underlie the diametric motor effects produced by SPNs in the two distinct spatial subdomains, we asked whether ^DMS^dSPNs have different connectivity with PV^+^ neurons and Npas1^+^ neurons compared to ^DLS^dSPN input. Similar to ^DLS^dSPNs that preferentially targeted Npas1^+^ neurons, ^DMS^dSPNs provided a stronger input to Npas1^+^ neurons compared to PV^+^ neurons, regardless of the stimulus intensity examined (**Figure 8a**). In addition, ^DMS^iSPNs had a stronger input to PV^+^ neurons than Npas1^+^ neurons (**Figure 8a**). DMS inputs shared a similar organization with DLS inputs (**Figure 8a**). A complete description of the input-output relationship for distinct DMS inputs is listed in **Table 4**.

To ascertain selective recruitment of striatal input from a unique spatial domain, all data so far were obtained from stimulation of SPNs locally either within the DMS or DLS. Although the slicing angle was chosen to maintain maximal striatopallidal connectivity, axons were inevitably severed. To ensure our observations were not confounded by axonal preservation, we recorded synaptic responses with full-field stimulation of SPN terminals in the GPe. As shown in **Figure 8a**, this approach yielded results that were highly concordant with that obtained from somatic stimulation. The spatial specificity of the stimulation was confirmed with sequential applications of tetrodotoxin (TTX, a voltage-gated sodium channel blocker) and 4-aminopyridine (4-AP, a voltage-gated potassium channel blocker) (Petreanu et al., 2009) (**Figure 8d**). Although IPSCs were abolished with TTX for both somatic (i.e., intrastriatal) and axonal (i.e., intrapallidal) stimulation, subsequent co-application of TTX with 4-AP selectively restored the IPSCs with terminal stimulation. These results thus confirmed that somatic stimulation, but not terminal stimulation, involved conducting events. In sum, dSPNs from both the DMS and DLS preferentially target Npas1^+^ neurons; whereas iSPNs from both spatial domains preferentially target PV^+^ neurons.

### The dSPN innervation to the GPe is selectively strengthened in a chronic PD model

Both circuit model and experimental data converge on the idea that an increased striatopallidal input contributes to the aberrant activity in the GPe in Parkinson’s disease (PD) (Albin et al., 1989; Wichmann and DeLong, 2003; Walters et al., 2007; Ballion et al., 2009; Kita and Kita, 2011a, b; Lemos et al., 2016; Ryan et al., 2018; McIver et al., 2019). Our previous study demonstrated that total striatal input in unidentified GPe neurons increased following chronic 6-OHDA lesion (Cui et al., 2016). However, the alterations of individual striatopallidal subcircuit (i.e., dSPN-PV^+^, dSPN-Npas1^+^, iSPN-PV^+^, and iSPN-Npas1^+^) were unknown. We therefore examined the alterations in the input-output relationship of these subcircuits in the chronic 6-OHDA lesion model of PD. ^DLS^dSPN-Npas1^+^ input was dramatically strengthened across all the stimulus intensities tested (IPSCs_max_: Npas1^+^_naïve_ = 213.8 ± 122.0 pA, *n* = 28 neurons; Npas1^+^_6-OHDA_ = 815.8 ± 530.5 pA, *n* = 15 neurons; *P* = 0.0017) (**Figure 8b**). In contrast, ^DLS^dSPN-PV^+^ input was not consistently altered across all stimulus intensities tested (IPSCs_max_: PV^+^_naïve_ = 38.5 ± 29.3 pA, *n* = 8 neurons; PV^+^_6-OHDA_ = 81.0 ± 76.7 pA, *n* = 13 neurons; *P* = 0.14) (**Figure 8b**). To our surprise, given a predicted increase in iSPN input with the classic circuit model (Albin et al., 1989; Wichmann and DeLong, 2003), neither ^DLS^iSPN-PV^+^ input nor ^DLS^iSPN-Npas1^+^ input was altered regardless of the stimulus location or stimulus intensity examined (IPSCs_max_: PV^+^_naïve_ = 1,740.7 ± 627.7 pA, *n* = 12 neurons; PV^+^_6-OHDA_ = 1,682.0 ± 820.4 pA, *n* = 18 neurons; *P* = 0.75; Npas1^+^_naïve_ = 76.9 ± 60.9 pA, *n* = 20 neurons; Npas1^+^_6-OHDA_ = 39.2 ± 22.2 pA, *n* = 11 neurons; *P* = 0.36) (**Figure 8a**). DMS inputs were similarly altered in the chronic 6-OHDA lesioned mice. ^DMS^dSPN-Npas1^+^ input was strengthened while no consistent change was detected for ^DMS^iSPN-PV^+^ input and ^DMS^iSPN-Npas1^+^ input (**Figure 8a**). As in naïve mice, our findings from 6-OHDA lesioned mice were consistent across the dStr subregions examined—dSPN inputs were biased toward Npas1^+^ neurons while iSPN inputs were biased toward PV^+^ neurons following chronic 6-OHDA lesion (**Figure 8a**). A full description of the input-output relationship for distinct striatal inputs is summarized in **Table 3 & 4**.

We hypothesized that the selective increase in the connection strength of the dSPN-Npas1^+^ input was a result of increased innervation. To test this idea, we measured the density of axons produced by dSPNs in naïve and 6-OHDA lesioned mice. To visualize dSPN axons, Cre-inducible ChR2-eYFP was expressed in *Drd1a*^Cre^ mice. We observed a dramatic increase in the dSPN axonal density in the GPe following 6-OHDA lesion (+2.64 ± 0.88 fold; *n*_naïve_ = 7 sections, 4 mice; *n*_6-OHDA_ = 12 sections, 6 mice; *P* = 0.00048) (**Figure 8e & g**), whereas only modest increases were observed in the dStr (+0.50 ± 0.22 fold; *n*_naïve_ = 12 sections, 4 mice; *n*_6-OHDA_ = 18 sections, 6 mice; *P* = 0.0050) and SNr (+0.57 ± 0.60 fold; *n*_naïve_ = 4 sections, 4 mice; *n*_6-OHDA_ = 6 sections, 6 mice; *P* = 0.48). The fold increase in the GPe was much higher than that in the dStr and SNr (GPe vs. dStr, *P* = 0.0087; GPe vs. SNr, *P* = 0.0087) (**Figure 8g**). As axonal density in the GPe reflects a sum of axonal arborizations and passage axons, synaptic contacts were quantified as an additional measure of innervation. Synaptic contacts were identified based on the overlap between eYFP^+^ and VGAT^+^ structures or the juxtaposition between eYFP^+^ and gephyrin^+^ elements (**Figure 8f & g**). The density of eYFP^+^-VGAT^+^ puncta in the GPe was greatly increased following 6-OHDA lesion (naïve = 3.9 ± 2.1 × 10^4^ count/mm^3^, *n* = 6 sections, 3 mice; 6-OHDA = 11.9 ± 3.0 × 10^4^ count/mm^3^, *n* = 10 sections, 5 mice; *P* = 0.0096). Additionally, a similar increase in the density of eYFP^+^-gephyrin^+^ puncta was found under the same conditions (naïve = 4.1 ± 1.5 × 10^4^ count/mm^3^, *n* = 6 sections, 3 mice; 6-OHDA = 19.3 ± 3.0 × 10^4^ count/mm^3^, *n* = 8 sections, 4 mice; *P* = 0.0020) (**Figure 8f & g**). As dSPN-Npas1^+^ input is the major input of dSPN projection, the increase in synaptic contacts from dSPNs explains the strengthening of the dSPN-Npas1^+^ input measured with *ex vivo* electrophysiology following chronic 6-OHDA lesion. In contrast, we did not find any alterations in release probability for the dSPN-Npas1^+^ input following 6-OHDA lesion, as indicated by the unaltered paired-pulse ratio (naïve = 1.1 ± 0.1, *n* = 10 neurons; 6-OHDA = 1.3 ± 0.2, *n* = 11 neurons; *P* = 0.20). In addition, no alterations in quantal amplitudes were observed following chronic 6-OHDA lesion (naïve = 86.1 ± 24.5 pA, *n* = 13 neurons; 6-OHDA = 92.8 ± 29.2 pA, *n* = 22 neurons; *P* = 0.39).

## Discussion

One of the central tenets of the basal ganglia model is the segregation of direct- and indirect-pathways, which selectively target the SNr and GPe, respectively. This canonical circuit model asserts that the two striatal output streams act as opponent pathways and are thought to promote and suppress motor output, respectively. Here, we showed that dSPNs (from both the DMS and DLS) target Npas1^+^ neurons in the GPe. Counter to the general belief that dSPNs are motor-promoting, selective stimulation of ^DLS^dSPNs led to motor suppression. As ^DLS^dSPN input to Npas1^+^ neurons was strengthened in a chronic model of PD, our results altogether suggest a previously unrecognized circuit regulation of motor function and dysfunction.

### iSPN-PV^+^ input is the principal component of the striatopallidal system

Here, we demonstrated that iSPN-PV^+^ input was the strongest connection among all striatopallidal inputs examined, namely, dSPN-PV^+^, dSPN-Npas1^+^, iSPN-PV^+^, and iSPN-Npas1^+^ inputs. Considering that PV^+^ neurons and Npas1^+^ neurons constitute 50% and 30% of all GPe neurons, respectively (Abdi et al., 2015; Dodson et al., 2015; Hernandez et al., 2015; Abecassis et al., 2020), iSPN-PV^+^ input should be regarded as the principal striatopallidal input.

Our finding that iSPN-PV^+^ input is stronger than iSPN-Npas1^+^ input is consistent with earlier anatomical studies showing that iSPNs form more synaptic contacts with PV^+^ neurons compared to PV^−^ neurons (Yuan et al., 2017). In addition, it is in line with recent electrophysiological studies that iSPNs strongly target Nkx2.1^+^ neurons (which are dominated by PV^+^ neurons) while providing minimal input to Foxp2^+^ neurons (which are a subset of Npas1^+^ neurons) (Yuan et al., 2017; Aristieta et al., 2020; Ketzef and Silberberg, 2020). Furthermore, we showed that the selective targeting of iSPN input to PV^+^ neurons was topographically conserved between spatial subdomains within the dStr—the DMS and the DLS.

At a circuit level, stimulation of iSPNs strongly suppresses firing of PV^+^ neurons, thus disinhibiting the targets of PV^+^ neurons, i.e., the STN and substantia nigra (Mastro et al., 2014; Hernandez et al., 2015; Saunders et al., 2016; Oh et al., 2017; Abecassis et al., 2020). As activity of PV^+^ neurons promotes movement (Cherian et al., 2020; Pamukcu et al., 2020), the selective targeting of PV^+^ neurons by iSPNs is in agreement with the movement suppressing role of ^DMS^iSPNs (Kravitz et al., 2010; Durieux et al., 2012). Conversely, we found stimulation of ^DLS^iSPNs promoted movement; this cannot be explained simply by the targeting properties of ^DLS^iSPNs. Further investigations are needed to determine the circuit mechanisms involved. In the present study, we did not find any alterations in the iSPN-PV^+^ input in the parkinsonian state. Given that iSPN-PV^+^ input is the principal striatopallidal input, this result is at odds with our previous observation that total striatopallidal input increases following chronic 6-OHDA lesion (Cui et al., 2016). As our prior study employed electrical stimulation, it is likely that subtle differences in the size and the waveform of the induced currents with optogenetic stimulation interfere with presynaptic regulation of release processes.

### The dSPN-Npas1^+^ input is functionally unique

In this study, we demonstrated that Npas1^+^ neurons are the principal target of dSPNs from both the DMS and DLS. Previous studies showed that dSPNs provide roughly half the number of boutons compared to iSPNs in the GPe (Kawaguchi et al., 1990; Fujiyama et al., 2011). Here, we found the relative number of boutons formed by iSPNs was seven times higher than that formed by dSPNs. Our electrophysiological data corroborate this anatomical finding; we found that the iSPN input to PV^+^ neurons was three to eight times greater than the dSPN input to Npas1^+^ neurons. The cell-targeting properties of the dSPN input was unexpected as earlier observations argue that the receptors for substance P are selectively expressed in PV^+^ neurons (Mizutani et al., 2017). These findings altogether suggest a target-specific innervation with transmitters released from a single cell type, i.e., GABA and substance P released from dSPNs preferentially inhibit Npas1^+^ neurons and excite PV^+^ neurons, respectively. Segregation of neurotransmitter release has also been found in other systems (Zhang et al., 2015; Lee et al., 2016; Granger et al., 2020). Given that Npas1^+^ neurons and PV^+^ neurons have opposite effects on motor control (Cherian et al., 2020; Pamukcu et al., 2020), the inhibition of Npas1^+^ neurons by GABA and excitation of PV^+^ neurons by substance P should work in concert to reinforce the same behavioral outcomes.

Despite the similar cell-targeting properties of ^DMS^dSPNs and ^DLS^dSPNs, they played dichotomous actions. We found that ^DLS^dSPNs exerted a movement-suppressing effect; this is opposite to the movement-promoting effect produced by ^DMS^dSPNs shown in this and other studies (Kravitz et al., 2010; Durieux et al., 2012; Freeze et al., 2013). Along this line, opposing regulation by ^DMS^dSPNs and ^DLS^dSPNs has also been found recently in regulating action sequence (Garr and Delamater, 2020). It is possible that differential recruitments of downstream pathways underlie the differences (**Figure 9**). As more PV^+^ neurons reside in the lateral GPe, stimulation of ^DLS^dSPNs should favor the engagement of the iSPN-PV^+^-SNr pathway through relay from Npas1^+^ neurons. On the contrary, as there are fewer PV^+^ neurons in the medial GPe, the effect of ^DMS^dSPNs is likely mediated through a disinhibition of the substantia nigra pars compacta via Npas1^+^ neurons (**Figure 9**).

**Figure 9.**
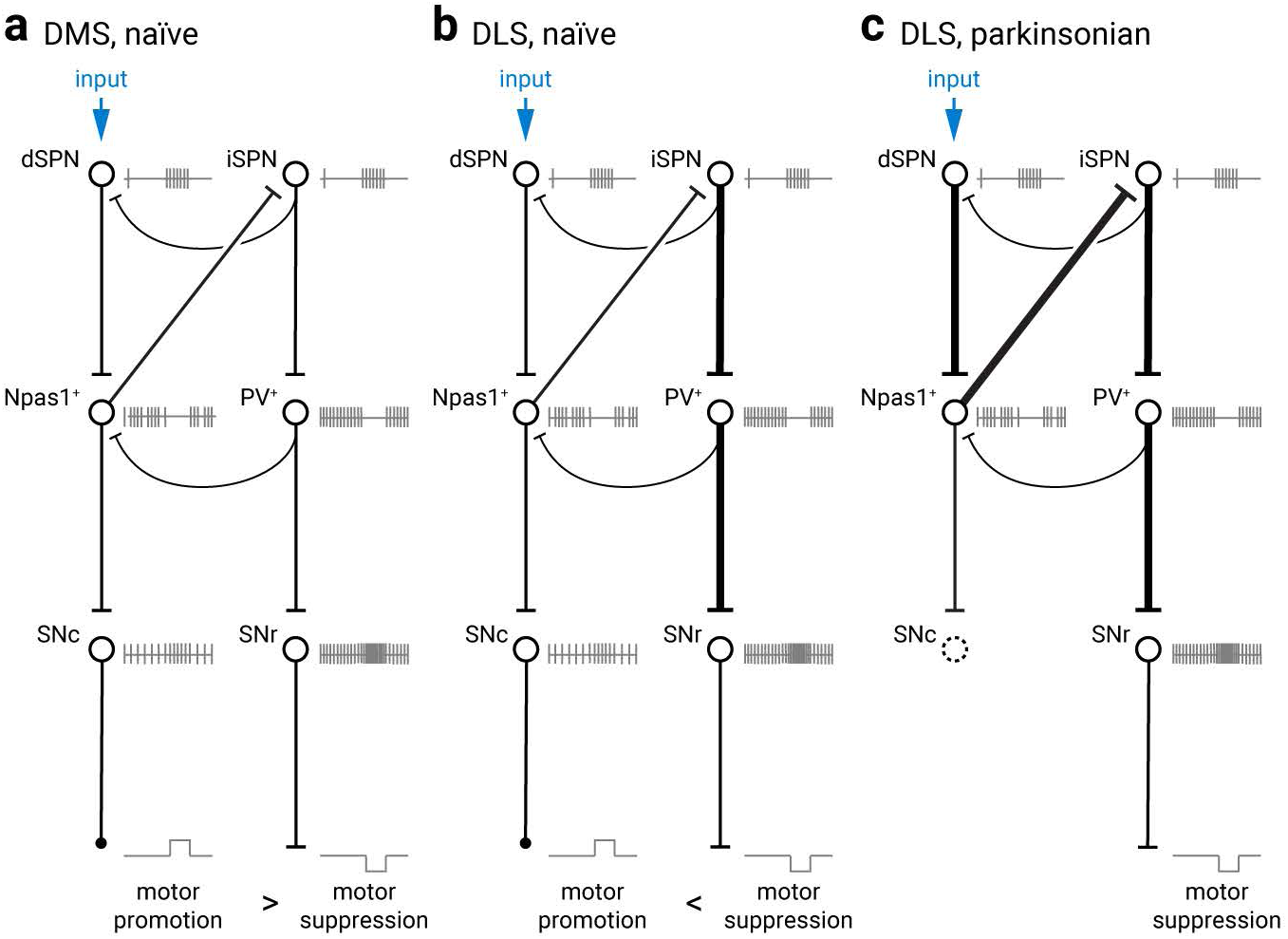
Schematic summary of proposed mechanisms for the observed motor responses. Npas1^+^ neurons and PV^+^ neurons in the GPe are the principal recipients of dSPN input and iSPN input, respectively. Npas1^+^ neurons in turn project to the dorsal striatum (dStr) and the substantia nigra pars compacta (SNc) (Abecassis et al., 2020; Cherian et al., 2020; Evans et al., 2020). PV^+^ neurons influence the output of substantia nigra pars reticulata (SNr) neurons via their direct projection and indirect projection through the subthalamic nucleus. Selective activation of dSPNs (e.g., with optogenetic stimulation) leads to either motor promotion or suppression depending on the balance of these two striatopallidal subcircuits. The coordination between the two subcircuits can be fine-tuned by local collaterals at the dStr, GPe, and SNr levels. Lateral inhibition between dSPNs and iSPNs is asymmetrical; iSPNs unidirectionally inhibit dSPNs (Taverna et al., 2008). PV^+^ neurons emit numerous local collaterals, thus providing additional crosstalk between the two striatopallidal subcircuits (Cherian et al., 2020). Local inhibitory circuits in the SNr have also been described; in particular, they can play important roles in regulating information transfer across the mediolateral axis (Brown et al., 2014). The same circuit motifs are conserved across the mediolateral extent of the basal ganglia. The diagrams serve as a working model that explains the behavioral effects observed in this study: DMS, naïve (**a**), DLS, naïve (**b**), and DLS, parkinsonian state (**c**). Stimulation of ^DMS^dSPNs has a strong motor-promotion effect as inhibition of Npas1^+^ neuron activity promotes the release of dopamine as a result of increased SNc neuron firing. In addition, as there are fewer PV^+^ neurons in the medial GPe (Kita, 1994; Hontanilla et al., 1998; Mastro et al., 2014; Hernandez et al., 2015; Abecassis et al., 2020), there is relatively weak engagement of the iSPN-PV^+^-SNr pathway. Furthermore, it likely strongly engages the canonical direct-pathway (from dSPNs to SNr) (not shown). Stimulation of ^DLS^dSPNs has a strong motor-suppression effect as there are more PV^+^ neurons in the lateral GPe. This favors engagement of the iSPN-PV^+^-SNr pathway and hence a net motor suppression. Following the loss of dopamine neurons, output from both dSPNs and Npas1^+^ neurons are both strengthened (Glajch et al., 2016). Despite the lack of changes in the connection strength of the iSPN-PV^+^ pathway, the strong disinhibition of iSPNs by Npas1^+^ neurons is expected to lead to strong disinhibition of the iSPN-PV^+^-SNr pathway, thus contributing to the hypokinetic symptoms of Parkinson’s disease.

The existence of parallel motor-suppressing striatopallidal pathways, i.e., ^DMS^iSPN-PV^+^ and ^DLS^dSPN-Npas1^+^, suggests a complex movement regulation by the dStr. Although stimulation of either pathway similarly decreased average speed, high-dimensional analyses revealed that the two pathways tune movement behavior in different patterns (**Figure 2**). It is conceivable that each functional subdomain within the dStr is equipped with both motor-suppressing and motor-promoting elements for the proper movement execution. Furthermore, the relative timing of recruitment of different subcircuits can play a role in shaping motor output (Yttri and Dudman, 2016).

### Limitations

First, our observations were limited to experiments with bulk manipulation (e.g., ChR2-mediated stimulation) of a relatively large population of neurons in the dStr. Recent studies suggest that motor command signals can be encoded by spatially confined neuronal ensembles in the dStr (Wiltschko et al., 2015; Barbera et al., 2016; Klaus et al., 2017; Girasole et al., 2018; Parker et al., 2018; Maltese et al., 2019; Sheng et al., 2019; Fobbs et al., 2020; Lee et al., 2020). While our data provided new insights into how motor patterns can be generated by striatal neuron subtypes across different spatial subdomains, close examination of the activity dynamics of these neurons and their synaptic partners in relation to movement will help clarify the precise circuit mechanisms involved.

Second, recent studies established the existence of two subclasses of Npas1^+^ neurons, i.e., Npas1^+^-Foxp2^+^ neurons and Npas1^+^-Nkx2.1^+^ (aka Npr3^+^) neurons that display distinct axonal projection patterns (Hernandez et al., 2015; Glajch et al., 2016; Saunders et al., 2018; Abecassis et al., 2020; Cherian et al., 2020). We do not currently know if dSPN input has differential impacts on these two neuron subclasses and their downstream targets. Finer scale circuit manipulations are necessary to fully understand the circuit mediating the effect of dSPNs.

### ^DLS^dSPN-Npas1^+^ pathway contributes to hypokinetic symptoms

In this study, we provided both anatomical and functional evidence that dSPN inputs to the GPe were strengthened following chronic 6-OHDA lesion. As activity of dopamine D2 receptors is known to positively regulate the density of dSPN innervation to the GPe, one would expect the inactivity of D2 receptors resulting from the 6-OHDA lesion to decrease this innervation. On the other hand, as iSPN activity promotes dSPN-GPe innervation (Cazorla et al., 2014), it is likely that the observed increase in dSPN-GPe innervation following chronic 6-OHDA lesion is a consequence of increased iSPN activity *in vivo*, as a result of the overactivity of excitatory inputs. This argument is supported by the observation that the activity of iSPNs (or presumptive iSPNs) are enhanced in animal models of PD (Calabresi et al., 1993; Schwarting and Huston, 1996; Kish et al., 1999; Pang et al., 2001; Tseng et al., 2001; Mallet et al., 2006; Kita and Kita, 2011a, b; Kovaleski et al., 2020) and that silencing of excitatory inputs selectively to iSPNs ameliorates motor deficits (Shields et al., 2017). As stimulation of ^DLS^dSPN-Npas1^+^ pathway suppresses movement, the increase in this input following chronic 6-OHDA lesion constitutes a novel mechanism underlying hypokinetic symptoms in this disease state.

## Acknowledgments

We would like to thank Lily Amadio, Saivasudha Chalasani, Alyssa Bebenek, Kris Shah, Moises Melesio, and Coby Dodelson for their assistance, Dr. Tracy Gertler, Cooper Chan, and Cassidy Chan for supporting the completion of the fourth manuscript from the Chan Lab during the COVID-19 pandemic. This work was supported by NIH R01 NS069777 (CSC), R01 MH112768 (CSC), R01 NS097901 (CSC), R01 MH109466 (CSC), R01 NS088528 (CSC), R01 MH107742 (BL), R01 MH108594 (BL), U01 MH114829 (BKL), R01 MH116176 (YK), and T32 AG020506 (AP).

## Author contributions

QC conceived the study. QC and XD conducted the electrophysiological measurements. QC, IYMC, and AP performed the behavioral studies. QC, XD, BLB, DG, and AN performed histological analysis. VL and BL performed the rabies tracing. UC and YK performed the two-photon tomography. DH and AN assisted with data analyses. QC and CSC wrote the manuscript. All authors reviewed and edited the manuscript. CSC designed, directed, and supervised the project.

## References

Abdi A, Mallet N, Mohamed FY, Sharott A, Dodson PD, Nakamura KC, Suri S, Avery SV, Larvin JT, Garas FN, Garas SN, Vinciati F, Morin S, Bezard E, Baufreton J, Magill PJ (2015) Prototypic and arkypallidal neurons in the dopamine-intact external globus pallidus. J Neurosci 35:6667–6688.

Abecassis ZA, Berceau BL, Win PH, Garcia D, Xenias HS, Cui Q, Pamukcu A, Cherian S, Hernandez VM, Chon U, Lim BK, Kim Y, Justice NJ, Awatramani R, Hooks BM, Gerfen CR, Boca SM, Chan CS (2020) Npas1(+)- Nkx2.1(+) Neurons Are an Integral Part of the Cortico-pallido-cortical Loop. J Neurosci 40:743–768.

Ade KK, Wan Y, Chen M, Gloss B, Calakos N (2011) An Improved BAC Transgenic Fluorescent Reporter Line for Sensitive and Specific Identification of Striatonigral Medium Spiny Neurons. Front Syst Neurosci 5:32.

Albin RL, Young AB, Penney JB (1989) The functional anatomy of basal ganglia disorders. Trends Neurosci 12:366–375.

Albin RL, Young AB, Penney JB (1995) The functional anatomy of disorders of the basal ganglia. Trends Neurosci 18:63–64.

Alegre-Cortés J, Sáez M, Montanari R, Reig R (2020) Medium spiny neurons activity reveals the discrete segregation of mouse dorsal striatum. bioRxiv:2020.2002.2021.959692.

Alexander GE, Crutcher MD (1990) Functional architecture of basal ganglia circuits: neural substrates of parallel processing. Trends Neurosci 13:266–271.

Aristieta A, Barresi M, Azizpour Lindi S, Barriere G, Courtand G, de la Crompe B, Guilhemsang L, Gauthier S, Fioramonti S, Baufreton J, Mallet NP (2020) A Disynaptic Circuit in the Globus Pallidus Controls Locomotion Inhibition. Curr Biol.

Balleine BW, O’Doherty JP (2010) Human and rodent homologies in action control: corticostriatal determinants of goal-directed and habitual action. Neuropsychopharmacology 35:48–69.

Ballion B, Frenois F, Zold CL, Chetrit J, Murer MG, Gonon F (2009) D2 receptor stimulation, but not D1, restores striatal equilibrium in a rat model of Parkinsonism. Neurobiol Dis 35:376–384.

Barbera G, Liang B, Zhang L, Gerfen CR, Culurciello E, Chen R, Li Y, Lin DT (2016) Spatially Compact Neural Clusters in the Dorsal Striatum Encode Locomotion Relevant Information. Neuron 92:202–213.

Bekkers JM, Clements JD (1999) Quantal amplitude and quantal variance of strontium-induced asynchronous EPSCs in rat dentate granule neurons. J Physiol 516 (Pt 1):227–248.

Bertino S, Basile GA, Bramanti A, Anastasi GP, Quartarone A, Milardi D, Cacciola A (2020) Spatially coherent and topographically organized pathways of the human globus pallidus. Hum Brain Mapp.

Boyden ES, Zhang F, Bamberg E, Nagel G, Deisseroth K (2005) Millisecond-timescale, genetically targeted optical control of neural activity. Nat Neurosci 8:1263–1268.

Brown J, Pan WX, Dudman JT (2014) The inhibitory microcircuit of the substantia nigra provides feedback gain control of the basal ganglia output. Elife 3:e02397.

Calabresi P, Mercuri NB, Sancesario G, Bernardi G (1993) Electrophysiology of dopamine-denervated striatal neurons. Implications for Parkinson’s disease. Brain 116 (Pt 2):433–452.

Cazorla M, de Carvalho FD, Chohan MO, Shegda M, Chuhma N, Rayport S, Ahmari SE, Moore H, Kellendonk C (2014) Dopamine D2 receptors regulate the anatomical and functional balance of basal ganglia circuitry. Neuron 81:153–164.

Chang HT, Wilson CJ, Kitai ST (1981) Single neostriatal efferent axons in the globus pallidus: a light and electron microscopic study. Science 213:915–918.

Cherian S, Cui Q, Pamukcu A, Chang IYM, Berceau BL, Xenias HS, Higgs MH, Rajamanickam S, Chen Y, Du X, Zhang Y, McMorrow H, Abecassis ZA, Boca SM, Justice NJ, Wilson CJ, Chan CS (2020) Dissociable Roles of Pallidal Neuron Subtypes in Regulating Motor Patterns. bioRxiv:2020.2008.2023.263053.

Chon U, Vanselow DJ, Cheng KC, Kim Y (2019) Enhanced and unified anatomical labeling for a common mouse brain atlas. Nat Commun 10:5067.

Conte WL, Kamishina H, Reep RL (2009a) Multiple neuroanatomical tract-tracing using fluorescent Alexa Fluor conjugates of cholera toxin subunit B in rats. Nat Protoc 4:1157–1166.

Conte WL, Kamishina H, Reep RL (2009b) The efficacy of the fluorescent conjugates of cholera toxin subunit B for multiple retrograde tract tracing in the central nervous system. Brain Struct Funct 213:367–373.

Costa RM (2011) A selectionist account of de novo action learning. Curr Opin Neurobiol 21:579–586.

Cowan WM, Powell TP (1966) Strio-pallidal projection in the monkey. J Neurol Neurosurg Psychiatry 29:426–439.

Cox J, Witten IB (2019) Striatal circuits for reward learning and decision-making. Nat Rev Neurosci 20:482–494.

Cui G, Jun SB, Jin X, Pham MD, Vogel SS, Lovinger DM, Costa RM (2013) Concurrent activation of striatal direct and indirect pathways during action initiation. Nature 494:238–242.

Cui Q, Pitt JE, Pamukcu A, Poulin JF, Mabrouk OS, Fiske MP, Fan IB, Augustine EC, Young KA, Kennedy RT, Awatramani R, Chan CS (2016) Blunted mGluR Activation Disinhibits Striatopallidal Transmission in Parkinsonian Mice. Cell Rep 17:2431–2444.

Darvas M, Palmiter RD (2009) Restriction of dopamine signaling to the dorsolateral striatum is sufficient for many cognitive behaviors. Proc Natl Acad Sci U S A 106:14664–14669.

Darvas M, Palmiter RD (2010) Restricting dopaminergic signaling to either dorsolateral or medial striatum facilitates cognition. J Neurosci 30:1158–1165.

DeLong MR, Wichmann T (2007) Circuits and circuit disorders of the basal ganglia. Arch Neurol 64:20–24.

Deniau JM, Menetrey A, Charpier S (1996) The lamellar organization of the rat substantia nigra pars reticulata: segregated patterns of striatal afferents and relationship to the topography of corticostriatal projections. Neuroscience 73:761–781.

Dodson PD, Larvin JT, Duffell JM, Garas FN, Doig NM, Kessaris N, Duguid IC, Bogacz R, Butt SJ, Magill PJ (2015) Distinct developmental origins manifest in the specialized encoding of movement by adult neurons of the external globus pallidus. Neuron 86:501–513.

Durieux PF, Schiffmann SN, de Kerchove d’Exaerde A (2012) Differential regulation of motor control and response to dopaminergic drugs by D1R and D2R neurons in distinct dorsal striatum subregions. EMBO J 31:640–653.

Evans RC, Twedell EL, Zhu M, Ascencio J, Zhang R, Khaliq ZM (2020) Functional Dissection of Basal Ganglia Inhibitory Inputs onto Substantia Nigra Dopaminergic Neurons. Cell Rep 32:108156.

Faget L, Zell V, Souter E, McPherson A, Ressler R, Gutierrez-Reed N, Yoo JH, Dulcis D, Hnasko TS (2018) Opponent control of behavioral reinforcement by inhibitory and excitatory projections from the ventral pallidum. Nat Commun 9:849.

Flaherty AW, Graybiel AM (1994) Input-output organization of the sensorimotor striatum in the squirrel monkey. J Neurosci 14:599–610.

Fobbs WC, Bariselli S, Licholai JA, Miyazaki NL, Matikainen-Ankney BA, Creed MC, Kravitz AV (2020) Continuous Representations of Speed by Striatal Medium Spiny Neurons. J Neurosci 40:1679–1688.

Foster NN et al. (2020) The mouse cortico-basal ganglia-thalamic network.2020.2010.2006.326876.

Freeze BS, Kravitz AV, Hammack N, Berke JD, Kreitzer AC (2013) Control of basal ganglia output by direct and indirect pathway projection neurons. J Neurosci 33:18531–18539.

Fujiyama F, Sohn J, Nakano T, Furuta T, Nakamura KC, Matsuda W, Kaneko T (2011) Exclusive and common targets of neostriatofugal projections of rat striosome neurons: a single neuron-tracing study using a viral vector. Eur J Neurosci 33:668–677.

Garr E, Delamater AR (2020) Chemogenetic inhibition in the dorsal striatum reveals regional specificity of direct and indirect pathway control of action sequencing. Neurobiol Learn Mem 169:107169.

Gerfen CR (1992) The neostriatal mosaic: multiple levels of compartmental organization in the basal ganglia. Annu Rev Neurosci 15:285–320.

Gerfen CR, Young WS, 3rd (1988) Distribution of striatonigral and striatopallidal peptidergic neurons in both patch and matrix compartments: an in situ hybridization histochemistry and fluorescent retrograde tracing study. Brain Res 460:161–167.

Gerfen CR, Surmeier DJ (2011) Modulation of striatal projection systems by dopamine. Annu Rev Neurosci 34:441–466.

Gerfen CR, Paletzki R, Heintz N (2013) GENSAT BAC cre-recombinase driver lines to study the functional organization of cerebral cortical and basal ganglia circuits. Neuron 80:1368–1383.

Gerfen CR, Engber TM, Mahan LC, Susel Z, Chase TN, Monsma FJ, Jr., Sibley DR (1990) D1 and D2 dopamine receptor-regulated gene expression of striatonigral and striatopallidal neurons. Science 250:1429–1432.

Girasole AE, Lum MY, Nathaniel D, Bair-Marshall CJ, Guenthner CJ, Luo L, Kreitzer AC, Nelson AB (2018) A Subpopulation of Striatal Neurons Mediates Levodopa-Induced Dyskinesia. Neuron 97:787–795 e786.

Glajch KE, Kelver DA, Hegeman DJ, Cui Q, Xenias HS, Augustine EC, Hernandez VM, Verma N, Huang TY, Luo M, Justice NJ, Chan CS (2016) Npas1+ Pallidal Neurons Target Striatal Projection Neurons. J Neurosci 36:5472–5488.

Gokce O, Stanley GM, Treutlein B, Neff NF, Camp JG, Malenka RC, Rothwell PE, Fuccillo MV, Sudhof TC, Quake SR (2016) Cellular Taxonomy of the Mouse Striatum as Revealed by Single-Cell RNA-Seq. Cell Rep 16:1126–1137.

Gong S, Doughty M, Harbaugh CR, Cummins A, Hatten ME, Heintz N, Gerfen CR (2007) Targeting Cre recombinase to specific neuron populations with bacterial artificial chromosome constructs. J Neurosci 27:9817–9823.

Granger AJ, Wang W, Robertson K, El-Rifai M, Zanello AF, Bistrong K, Saunders A, Chow BW, Nunez V, Turrero Garcia M, Harwell CC, Gu C, Sabatini BL (2020) Cortical ChAT(+) neurons co-transmit acetylcholine and GABA in a target- and brain-region-specific manner. Elife 9.

Graybiel AM (2008) Habits, rituals, and the evaluative brain. Annu Rev Neurosci 31:359–387.

Harris JA, Hirokawa KE, Sorensen SA, Gu H, Mills M, Ng LL, Bohn P, Mortrud M, Ouellette B, Kidney J, Smith KA, Dang C, Sunkin S, Bernard A, Oh SW, Madisen L, Zeng H (2014) Anatomical characterization of Cre driver mice for neural circuit mapping and manipulation. Front Neural Circuits 8:76.

Hedreen JC, DeLong MR (1991) Organization of striatopallidal, striatonigral, and nigrostriatal projections in the macaque. J Comp Neurol 304:569–595.

Hegeman DJ, Hong ES, Hernandez VM, Chan CS (2016) The external globus pallidus: progress and perspectives. Eur J Neurosci 43:1239–1265.

Heiman M, Schaefer A, Gong S, Peterson JD, Day M, Ramsey KE, Suarez-Farinas M, Schwarz C, Stephan DA, Surmeier DJ, Greengard P, Heintz N (2008) A translational profiling approach for the molecular characterization of CNS cell types. Cell 135:738–748.

Hernandez VM, Hegeman DJ, Cui Q, Kelver DA, Fiske MP, Glajch KE, Pitt JE, Huang TY, Justice NJ, Chan CS (2015) Parvalbumin+ Neurons and Npas1+ Neurons Are Distinct Neuron Classes in the Mouse External Globus Pallidus. J Neurosci 35:11830–11847.

Hintiryan H, Foster NN, Bowman I, Bay M, Song MY, Gou L, Yamashita S, Bienkowski MS, Zingg B, Zhu M, Yang XW, Shih JC, Toga AW, Dong HW (2016) The mouse cortico-striatal projectome. Nat Neurosci 19:1100–1114.

Hippenmeyer S, Vrieseling E, Sigrist M, Portmann T, Laengle C, Ladle DR, Arber S (2005) A developmental switch in the response of DRG neurons to ETS transcription factor signaling. PLoS Biol 3:e159.

Hontanilla B, Parent A, de las Heras S, Gimenez-Amaya JM (1998) Distribution of calbindin D-28k and parvalbumin neurons and fibers in the rat basal ganglia. Brain Res Bull 47:107–116.

Hooks BM, Papale AE, Paletzki RF, Feroze MW, Eastwood BS, Couey JJ, Winnubst J, Chandrashekar J, Gerfen CR (2018) Topographic precision in sensory and motor corticostriatal projections varies across cell type and cortical area. Nat Commun 9:3549.

Hunnicutt BJ, Jongbloets BC, Birdsong WT, Gertz KJ, Zhong H, Mao T (2016) A comprehensive excitatory input map of the striatum reveals novel functional organization. Elife 5.

Hunt AJ, Jr., Dasgupta R, Rajamanickam S, Jiang Z, Beierlein M, Chan CS, Justice NJ (2018) Paraventricular hypothalamic and amygdalar CRF neurons synapse in the external globus pallidus. Brain Struct Funct 223:2685–2698.

Kaiser T, Ting JT, Monteiro P, Feng G (2016) Transgenic labeling of parvalbumin-expressing neurons with tdTomato. Neuroscience 321:236–245.

Kawaguchi Y, Wilson CJ, Emson PC (1990) Projection subtypes of rat neostriatal matrix cells revealed by intracellular injection of biocytin. J Neurosci 10:3421–3438.

Ketzef M, Silberberg G (2020) Differential Synaptic Input to External Globus Pallidus Neuronal Subpopulations In Vivo. Neuron.

Kim J, Alger BE (2001) Random response fluctuations lead to spurious paired-pulse facilitation. J Neurosci 21:9608–9618.

Kim Y, Yang GR, Pradhan K, Venkataraju KU, Bota M, Garcia Del Molino LC, Fitzgerald G, Ram K, He M, Levine JM, Mitra P, Huang ZJ, Wang XJ, Osten P (2017) Brain-wide Maps Reveal Stereotyped Cell-Type-Based Cortical Architecture and Subcortical Sexual Dimorphism. Cell 171:456–469 e422.

Kish LJ, Palmer MR, Gerhardt GA (1999) Multiple single-unit recordings in the striatum of freely moving animals: effects of apomorphine and D-amphetamine in normal and unilateral 6-hydroxydopamine-lesioned rats. Brain Res 833:58–70.

Kita H (1994) Parvalbumin-immunopositive neurons in rat globus pallidus: a light and electron microscopic study. Brain Res 657:31–41.

Kita H (2007) Globus pallidus external segment. Prog Brain Res 160:111–133.

Kita H, Kita T (2011a) Role of Striatum in the Pause and Burst Generation in the Globus Pallidus of 6-OHDA-Treated Rats. Front Syst Neurosci 5:42.

Kita H, Kita T (2011b) Cortical stimulation evokes abnormal responses in the dopamine-depleted rat basal ganglia. J Neurosci 31:10311–10322.

Klaus A, Martins GJ, Paixao VB, Zhou P, Paninski L, Costa RM (2017) The Spatiotemporal Organization of the Striatum Encodes Action Space. Neuron 95:1171–1180 e1177.

Knowland D, Lilascharoen V, Pacia CP, Shin S, Wang EH, Lim BK (2017) Distinct Ventral Pallidal Neural Populations Mediate Separate Symptoms of Depression. Cell 170:284–297 e218.

Kovaleski RF, Callahan JW, Chazalon M, Wokosin DL, Baufreton J, Bevan MD (2020) Dysregulation of external globus pallidus-subthalamic nucleus network dynamics in parkinsonian mice during cortical slow-wave activity and activation. J Physiol 598:1897–1927.

Kravitz AV, Tye LD, Kreitzer AC (2012) Distinct roles for direct and indirect pathway striatal neurons in reinforcement. Nat Neurosci 15:816–818.

Kravitz AV, Freeze BS, Parker PR, Kay K, Thwin MT, Deisseroth K, Kreitzer AC (2010) Regulation of parkinsonian motor behaviours by optogenetic control of basal ganglia circuitry. Nature 466:622–626.

Krzywinski M, Altman N (2014) Visualizing samples with box plots. Nat Methods 11:119–120.

Lee J, Wang W, Sabatini BL (2020) Anatomically segregated basal ganglia pathways allow parallel behavioral modulation. Nat Neurosci 23:1388–1398.

Lee S, Zhang Y, Chen M, Zhou ZJ (2016) Segregated Glycine-Glutamate Co-transmission from vGluT3 Amacrine Cells to Contrast-Suppressed and Contrast-Enhanced Retinal Circuits. Neuron 90:27–34.

Lemos JC, Friend DM, Kaplan AR, Shin JH, Rubinstein M, Kravitz AV, Alvarez VA (2016) Enhanced GABA Transmission Drives Bradykinesia Following Loss of Dopamine D2 Receptor Signaling. Neuron 90:824–838.

Levesque M, Parent A (2005) The striatofugal fiber system in primates: a reevaluation of its organization based on single-axon tracing studies. Proc Natl Acad Sci U S A 102:11888–11893.

Leys C, Ley C, Klein O, Bernard P, Licata L (2013) Detecting outliers: Do not use standard deviation around the mean, use absolute deviation around the median. Journal of Experimental Social Psychology 49:764–766.

Lobo MK, Karsten SL, Gray M, Geschwind DH, Yang XW (2006) FACS-array profiling of striatal projection neuron subtypes in juvenile and adult mouse brains. Nat Neurosci 9:443–452.

Lobo MK, Covington HE, 3rd, Chaudhury D, Friedman AK, Sun H, Damez-Werno D, Dietz DM, Zaman S, Koo JW, Kennedy PJ, Mouzon E, Mogri M, Neve RL, Deisseroth K, Han MH, Nestler EJ (2010) Cell type-specific loss of BDNF signaling mimics optogenetic control of cocaine reward. Science 330:385–390.

Madisen L, Zwingman TA, Sunkin SM, Oh SW, Zariwala HA, Gu H, Ng LL, Palmiter RD, Hawrylycz MJ, Jones AR, Lein ES, Zeng H (2010) A robust and high-throughput Cre reporting and characterization system for the whole mouse brain. Nat Neurosci 13:133–140.

Madisen L et al. (2015) Transgenic mice for intersectional targeting of neural sensors and effectors with high specificity and performance. Neuron 85:942–958.

Mahn M, Gibor L, Patil P, Cohen-Kashi Malina K, Oring S, Printz Y, Levy R, Lampl I, Yizhar O (2018) High-efficiency optogenetic silencing with soma-targeted anion-conducting channelrhodopsins. Nat Commun 9:4125.

Mallet N, Ballion B, Le Moine C, Gonon F (2006) Cortical inputs and GABA interneurons imbalance projection neurons in the striatum of parkinsonian rats. J Neurosci 26:3875–3884.

Mallet N, Schmidt R, Leventhal D, Chen F, Amer N, Boraud T, Berke JD (2016) Arkypallidal Cells Send a Stop Signal to Striatum. Neuron 89:308–316.

Maltese M, March JR, Bashaw AG, Tritsch NX (2019) Dopamine modulates the size of striatal projection neuron ensembles.865006.

Malvaez M, Wassum KM (2018) Regulation of habit formation in the dorsal striatum. Curr Opin Behav Sci 20:67–74.

Markowitz JE, Gillis WF, Beron CC, Neufeld SQ, Robertson K, Bhagat ND, Peterson RE, Peterson E, Hyun M, Linderman SW, Sabatini BL, Datta SR (2018) The Striatum Organizes 3D Behavior via Moment-to-Moment Action Selection. Cell 174:44–58 e17.

Martin A, Calvigioni D, Tzortzi O, Fuzik J, Warnberg E, Meletis K (2019) A Spatiomolecular Map of the Striatum. Cell Rep 29:4320–4333 e4325.

Mastro KJ, Bouchard RS, Holt HA, Gittis AH (2014) Transgenic mouse lines subdivide external segment of the globus pallidus (GPe) neurons and reveal distinct GPe output pathways. J Neurosci 34:2087–2099.

Mathis A, Yüksekgönül M, Rogers B, Bethge M, Mathis MW (2019) Pretraining Boosts Out-of-domain Robustness for Pose Estimation. arXiv:1909.11229.

Mathis A, Mamidanna P, Cury KM, Abe T, Murthy VN, Mathis MW, Bethge M (2018) DeepLabCut: markerless pose estimation of user-defined body parts with deep learning. Nat Neurosci 21:1281–1289.

McGarry LM, Carter AG (2017) Prefrontal Cortex Drives Distinct Projection Neurons in the Basolateral Amygdala. Cell Rep 21:1426–1433.

McGeorge AJ, Faull RL (1989) The organization of the projection from the cerebral cortex to the striatum in the rat. Neuroscience 29:503–537.

McIver EL, Atherton JF, Chu HY, Cosgrove KE, Kondapalli J, Wokosin D, Surmeier DJ, Bevan MD (2019) Maladaptive Downregulation of Autonomous Subthalamic Nucleus Activity following the Loss of Midbrain Dopamine Neurons. Cell Rep 28:992–1002 e1004.

Metsalu T, Vilo J (2015) ClustVis: a web tool for visualizing clustering of multivariate data using Principal Component Analysis and heatmap. Nucleic Acids Res 43:W566–570.

Mink JW (1996) The basal ganglia: focused selection and inhibition of competing motor programs. Prog Neurobiol 50:381–425.

Mizutani K, Takahashi S, Okamoto S, Karube F, Fujiyama F (2017) Substance P effects exclusively on prototypic neurons in mouse globus pallidus. Brain Struct Funct 222:4089–4110.

Nambu A (2011) Somatotopic organization of the primate Basal Ganglia. Front Neuroanat 5:26.

Nath T, Mathis A, Chen AC, Patel A, Bethge M, Mathis MW (2019) Using DeepLabCut for 3D markerless pose estimation across species and behaviors. Nat Protoc 14:2152–2176.

Nuzzo RL (2016) The Box Plots Alternative for Visualizing Quantitative Data. PM R 8:268–272.

O’Hare JK, Ade KK, Sukharnikova T, Van Hooser SD, Palmeri ML, Yin HH, Calakos N (2016) Pathway-Specific Striatal Substrates for Habitual Behavior. Neuron 89:472–479.

Oh SW et al. (2014) A mesoscale connectome of the mouse brain. Nature 508:207–214.

Oh YM, Karube F, Takahashi S, Kobayashi K, Takada M, Uchigashima M, Watanabe M, Nishizawa K, Kobayashi K, Fujiyama F (2017) Using a novel PV-Cre rat model to characterize pallidonigral cells and their terminations. Brain Struct Funct 222:2359–2378.

Okamoto S, Sohn J, Tanaka T, Takahashi M, Ishida Y, Yamauchi K, Koike M, Fujiyama F, Hioki H (2020) Overlapping Projections of Neighboring Direct and Indirect Pathway Neostriatal Neurons to Globus Pallidus External Segment. iScience:101409.

Ortiz C, Navarro JF, Jurek A, Martin A, Lundeberg J, Meletis K (2020) Molecular atlas of the adult mouse brain. Sci Adv 6:eabb3446.

Owen SF, Liu MH, Kreitzer AC (2019) Thermal constraints on in vivo optogenetic manipulations. Nat Neurosci 22:1061–1065.

Pamukcu A, Cui Q, Xenias HS, Berceau BL, Augustine EC, Fan I, Chalasani S, Hantman AW, Lerner TN, Boca SM, Chan CS (2020) Parvalbumin(+) and Npas1(+) Pallidal Neurons Have Distinct Circuit Topology and Function. J Neurosci 40:7855–7876.

Pan WX, Mao T, Dudman JT (2010) Inputs to the dorsal striatum of the mouse reflect the parallel circuit architecture of the forebrain. Front Neuroanat 4:147.

Pang Z, Ling GY, Gajendiran M, Xu ZC (2001) Enhanced excitatory synaptic transmission in spiny neurons of rat striatum after unilateral dopamine denervation. Neurosci Lett 308:201–205.

Park J, Coddington LT, Dudman JT (2020) Basal Ganglia Circuits for Action Specification. Annu Rev Neurosci 43:485–507.

Parker JG, Marshall JD, Ahanonu B, Wu YW, Kim TH, Grewe BF, Zhang Y, Li JZ, Ding JB, Ehlers MD, Schnitzer MJ (2018) Diametric neural ensemble dynamics in parkinsonian and dyskinetic states. Nature 557:177–182.

Petreanu L, Mao T, Sternson SM, Svoboda K (2009) The subcellular organization of neocortical excitatory connections. Nature 457:1142–1145.

Poulin JF, Caronia G, Hofer C, Cui Q, Helm B, Ramakrishnan C, Chan CS, Dombeck DA, Deisseroth K, Awatramani R (2018) Mapping projections of molecularly defined dopamine neuron subtypes using intersectional genetic approaches. Nat Neurosci 21:1260–1271.

Redgrave P, Rodriguez M, Smith Y, Rodriguez-Oroz MC, Lehericy S, Bergman H, Agid Y, DeLong MR, Obeso JA (2010) Goal-directed and habitual control in the basal ganglia: implications for Parkinson’s disease. Nat Rev Neurosci 11:760–772.

Romanelli P, Esposito V, Schaal DW, Heit G (2005) Somatotopy in the basal ganglia: experimental and clinical evidence for segregated sensorimotor channels. Brain Res Brain Res Rev 48:112–128.

Rothwell PE, Fuccillo MV, Maxeiner S, Hayton SJ, Gokce O, Lim BK, Fowler SC, Malenka RC, Sudhof TC (2014) Autism-associated neuroligin-3 mutations commonly impair striatal circuits to boost repetitive behaviors. Cell 158:198–212.

Ryan MB, Bair-Marshall C, Nelson AB (2018) Aberrant Striatal Activity in Parkinsonism and Levodopa-Induced Dyskinesia. Cell Rep 23:3438–3446 e3435.

Sandler M, Howard A, Zhu M, Zhmoginov A, Chen LC (2019) MobileNetV2: Inverted Residuals and Linear Bottlenecks. arXiv:1801.04381.

Saunders A, Huang KW, Sabatini BL (2016) Globus Pallidus Externus Neurons Expressing parvalbumin Interconnect the Subthalamic Nucleus and Striatal Interneurons. PLoS One 11:e0149798.

Saunders A, Macosko EZ, Wysoker A, Goldman M, Krienen FM, de Rivera H, Bien E, Baum M, Bortolin L, Wang S, Goeva A, Nemesh J, Kamitaki N, Brumbaugh S, Kulp D, McCarroll SA (2018) Molecular Diversity and Specializations among the Cells of the Adult Mouse Brain. Cell 174:1015–1030 e1016.

Schindelin J, Arganda-Carreras I, Frise E, Kaynig V, Longair M, Pietzsch T, Preibisch S, Rueden C, Saalfeld S, Schmid B, Tinevez JY, White DJ, Hartenstein V, Eliceiri K, Tomancak P, Cardona A (2012) Fiji: an open-source platform for biological-image analysis. Nat Methods 9:676–682.

Schwarting RK, Huston JP (1996) The unilateral 6-hydroxydopamine lesion model in behavioral brain research. Analysis of functional deficits, recovery and treatments. Prog Neurobiol 50:275–331.

Sheng MJ, Lu D, Shen ZM, Poo MM (2019) Emergence of stable striatal D1R and D2R neuronal ensembles with distinct firing sequence during motor learning. Proc Natl Acad Sci U S A 116:11038–11047.

Shields BC, Kahuno E, Kim C, Apostolides PF, Brown J, Lindo S, Mensh BD, Dudman JT, Lavis LD, Tadross MR (2017) Deconstructing behavioral neuropharmacology with cellular specificity. Science 356.

Shin S, Pribiag H, Lilascharoen V, Knowland D, Wang XY, Lim BK (2018) Drd3 Signaling in the Lateral Septum Mediates Early Life Stress-Induced Social Dysfunction. Neuron 97:195–208 e196.

Smith Y, Raju DV, Pare JF, Sidibe M (2004) The thalamostriatal system: a highly specific network of the basal ganglia circuitry. Trends Neurosci 27:520–527.

Streit M, Gehlenborg N (2014) Bar charts and box plots. Nat Methods 11:117.

Szabo J (1962) Topical Distribution of the Striatal Efferents in the Monkey. Exp Neurol 5:21–36.

Taverna S, Ilijic E, Surmeier DJ (2008) Recurrent collateral connections of striatal medium spiny neurons are disrupted in models of Parkinson’s disease. J Neurosci 28:5504–5512.

Tseng KY, Kasanetz F, Kargieman L, Riquelme LA, Murer MG (2001) Cortical slow oscillatory activity is reflected in the membrane potential and spike trains of striatal neurons in rats with chronic nigrostriatal lesions. J Neurosci 21:6430–6439.

Turner RS, Desmurget M (2010) Basal ganglia contributions to motor control: a vigorous tutor. Curr Opin Neurobiol 20:704–716.

Walters JR, Hu D, Itoga CA, Parr-Brownlie LC, Bergstrom DA (2007) Phase relationships support a role for coordinated activity in the indirect pathway in organizing slow oscillations in basal ganglia output after loss of dopamine. Neuroscience 144:762–776.

Weglage M, Wärnberg E, Lazaridis I, Tzortzi O, Meletis K (2020) Complete representation of action space and value in all striatal pathways. bioRxiv:2020.2003.2029.983825.

Wichmann T, DeLong MR (2003) Pathophysiology of Parkinson’s disease: the MPTP primate model of the human disorder. Ann N Y Acad Sci 991:199–213.

Wilson CJ, Phelan KD (1982) Dual topographic representation of neostriatum in the globus pallidus of rats. Brain Res 243:354–359.

Wiltschko AB, Johnson MJ, Iurilli G, Peterson RE, Katon JM, Pashkovski SL, Abraira VE, Adams RP, Datta SR (2015) Mapping Sub-Second Structure in Mouse Behavior. Neuron 88:1121–1135.

Wu Y, Richard S, Parent A (2000) The organization of the striatal output system: a single-cell juxtacellular labeling study in the rat. Neurosci Res 38:49–62.

Xu-Friedman MA, Regehr WG (1999) Presynaptic strontium dynamics and synaptic transmission. Biophys J 76:2029–2042.

Xu-Friedman MA, Regehr WG (2000) Probing fundamental aspects of synaptic transmission with strontium. J Neurosci 20:4414–4422.

Yin HH, Knowlton BJ (2006) The role of the basal ganglia in habit formation. Nat Rev Neurosci 7:464–476.

Yttri EA, Dudman JT (2016) Opponent and bidirectional control of movement velocity in the basal ganglia. Nature 533:402–406.

Yuan XS, Wang L, Dong H, Qu WM, Yang SR, Cherasse Y, Lazarus M, Schiffmann SN, d’Exaerde AK, Li RX, Huang ZL (2017) Striatal adenosine A2A receptor neurons control active-period sleep via parvalbumin neurons in external globus pallidus. Elife 6.

Zhang S, Qi J, Li X, Wang HL, Britt JP, Hoffman AF, Bonci A, Lupica CR, Morales M (2015) Dopaminergic and glutamatergic microdomains in a subset of rodent mesoaccumbens axons. Nat Neurosci 18:386–392.

Znamenskiy P, Zador AM (2013) Corticostriatal neurons in auditory cortex drive decisions during auditory discrimination. Nature 497:482–485.

